# Integrating multi-omics data and machine learning in single-step GBLUP reaction norm models to improve prediction across untested environments in lodgepole pine

**DOI:** 10.64898/2026.11.16.767158

**Authors:** Eduardo P. Cappa, Charles Chen, Andy Benowicz, Barb R. Thomas, Yousry A. El-Kassaby

**Affiliations:** Instituto Nacional de Tecnología Agropecuaria (INTA), Instituto de Recursos Biológicos, Centro de Investigación en Recursos Naturales, De Los Reseros y Dr. Nicolás Repetto s/n, 1686, Hurlingham, Buenos Aires, Argentina; Consejo Nacional de Investigaciones Científicas y Técnicas (CONICET), Buenos Aires, Argentina; Department of Biochemistry and Molecular Biology, Oklahoma State University, Stillwater, Oklahoma 74078, USA; Department of Renewable Resources, University of Alberta, 442 Earth Sciences Bldg., Edmonton, Alberta, T6G 2E3, Canada; Forest Stewardship and Trade Branch, Alberta Forestry and Parks, Edmonton, Alberta, T6H 5T6, Canada; Department of Forest and Conservation Sciences, Faculty of Forestry, University of British Columbia, Vancouver, British Columbia, V6T 1Z4, Canada

**Keywords:** multi-omics integration, genotype-by-environment interaction, reaction norm models, nonlinear kernels, genomic prediction

## Abstract

The effectiveness of genomic prediction in forest trees is often constrained by genotype-by-environment (G×E) interactions, particularly when predicting performance in environments not represented in the training population. We evaluated the integration of genomic, enviromic (climatic and soil), and metabolomic (targeted monoterpene) information within multi-environment single-step genomic best linear unbiased prediction (ssGBLUP) reaction norm models to improve prediction in unobserved environments. Models incorporated environmental covariates (W) and genotype-specific responses using linear and nonlinear kernels, including Gaussian and Arc-Cosine formulations. The study examined 12 traits measured on 11,124 trees across four lodgepole pine progeny trials, with 1,490 trees genotyped with 25,099 SNPs. Predictive performance was assessed using leave-one-environment-out cross-validation. Additive genetic correlations among environments revealed substantial G×E for early-growth traits and drought-related decline, with correlations frequently below 0.65. Inclusion of environmental covariates reduced variance attributed to site effects, indicating that these descriptors captured environmental heterogeneity. Incorporating G×E and genotype-by-environmental covariate (G×W) interactions improved model performance, increasing explained phenotypic variance by up to 29.58% and reducing residual variance by up to 3.13% relative to baseline models. Across traits, G×E explained more phenotypic variation than G×W, contributing on average 2.31% and 0.60%, respectively. Predictive ability was high across models (0.688-0.701), with gains of up to 7.85% when nonlinear kernels were combined with metabolomic covariates, particularly for traits exhibiting stronger G×E. Monoterpene profiles yielded predictive performance comparable to climate and soil descriptors, highlighting the potential of integrating genomic, enviromic, and metabolomic information to improve prediction across environments and support climate-resilient forest tree breeding and deployment.

## Introduction

Genomic prediction (GP) has emerged as a powerful tool to increase selection accuracy and accelerate breeding cycles by incorporating genome-wide marker information into breeding programs (Meuwissen *et al*., 2001). In forest tree species, GP is commonly implemented using genomic best linear unbiased prediction (GBLUP), which relies on genomic relationship matrices derived from SNP markers (VanRaden, 2008), and has been successfully applied across a range of species (Grattapaglia, 2022; Isik, 2022). More recently, single-step GBLUP (ssGBLUP) approaches have gained attention because they jointly integrate pedigree, phenotypic, and genomic information within a unified framework (Legarra *et al*., 2009; Misztal *et al*., 2009; Aguilar *et al*., 2010; Christensen and Lund, 2010). This feature is particularly attractive for forest tree breeding programs, where only a subset of individuals is typically genotyped (e.g., (Ratcliffe *et al*., 2017; Cappa *et al*., 2019; Callister *et al*., 2021)).

Despite these advances, the predictive ability of GP models can be compromised when genotype-by-environment (G×E) effects are strong (Resende *et al*., 2012a; Beaulieu *et al*., 2014; Grattapaglia, 2014; Duarte *et al*., 2024), as marker effects may not be consistently expressed across environments (Cappa *et al*., 2025). Multi-environment models have therefore been developed to account for G×E, either by modeling genetic correlations among environments or through reaction norm approaches. Traditional multi-trait models treat the same trait measured across environments as distinct traits and quantify G×E through genetic correlations among sites (Falconer and Mackay, 1996). Although effective in forest trees (e.g., (Li *et al*., 2007; Belaber *et al*., 2018; Cappa, Chen, *et al*., 2022)), these models become increasingly complex as the number of environments increases, even when more parsimonious structures such as factor-analytic models are used (Cullis *et al*., 2014). Moreover, they cannot predict performance in untested environments (Smith *et al*., 2015).

Reaction norm models provide an alternative framework by incorporating environmental covariates (ECs) to characterize environmental variation. Within this context, the enviromics framework uses large-scale environmental data (Costa-Neto and Fritsche-Neto, 2021) to construct environmental relationship matrices and model genotype-specific responses across environmental gradients (Jarquín *et al*., 2014). Building on this concept, (Pérez-Rodríguez *et al*., 2017) incorporated pedigree and genomic information within a ssGBLUP reaction norm model to predict wheat performance across international environments, demonstrating the potential of ssGBLUP models integrating G×E interactions for prediction in untested environments. While this formulation, using pedigree or genomic information, or both, has been widely applied in agricultural crops (Pérez-Rodríguez *et al*., 2015; Sukumaran *et al*., 2017; Morais Júnior *et al*., 2018), apple (Jung *et al*., 2025), alfalfa (Filho *et al*., 2023) and animal breeding (Ribeiro *et al*., 2015; Tiezzi *et al*., 2017; McWhorter *et al*., 2023; Bussiman *et al*., 2025), its application in forest trees remains limited (Ratcliffe *et al*., 2019; Oliveira Silva da Costa *et al*., 2025).

The increasing availability of high-resolution environmental datasets has further expanded enviromics approaches. Climate information from sources such as ClimateNA (https://climatena.ca/) and NASA POWER (https://power.larc.nasa.gov/), together with edaphic information from the Global Soil Information System (GLOSIS, https://data.apps.fao.org/glosis/), can be used to construct environmental similarity matrices (Costa-Neto *et al*., 2021; Fritsche-Neto, 2024). Beyond these traditional descriptors, emerging -omics technologies offer new opportunities to characterize biological responses to environmental variation and improve the modeling of G×E interactions.

Among the available -omics layers, metabolomics is particularly promising because metabolites are downstream products of gene expression and are therefore closely associated with phenotypic variation (Guijas *et al*., 2018). Metabolite profiles integrate both genetic and environmental influences, making them valuable biomarkers of environmental adaptation and G×E interactions (Wang et al., 2023). Unlike climatic and soil descriptors, metabolomic variables may also capture integrated physiological responses associated with environmental adaptation and stress (Yoshioka *et al*., 2026). Targeted metabolomics approaches, which quantify specific metabolites or metabolic pathways (Patti *et al*., 2012), provide a powerful framework to investigate biologically meaningful processes underlying plant responses. In conifers, for instance, constitutive monoterpene profiles have been linked to resistance against insect-pathogen complexes and may provide useful predictors for defense-related traits (Ullah *et al*., 2021). Consequently, metabolomic variables may serve not only as intermediate phenotypes but also as biologically informed descriptors of environmental similarity within reaction norm frameworks.

Multi-omics prediction integrates multiple sources of biological information to capture complementary processes underlying complex traits (see Figure 1 in (Robinson et al., 2025)). By incorporating intermediate phenotypes and regulatory mechanisms linking genotype to phenotype, these approaches can improve both predictive performance and biological interpretability (Xu *et al*., 2022; Montesinos-López *et al*., 2025; Yoshioka *et al*., 2026). However, identifying which omics variables are biologically meaningful and effectively improve prediction accuracy remains largely unexplored, particularly in the context of G×E interactions (Yoshioka *et al*., 2026).

**Figure 1.**
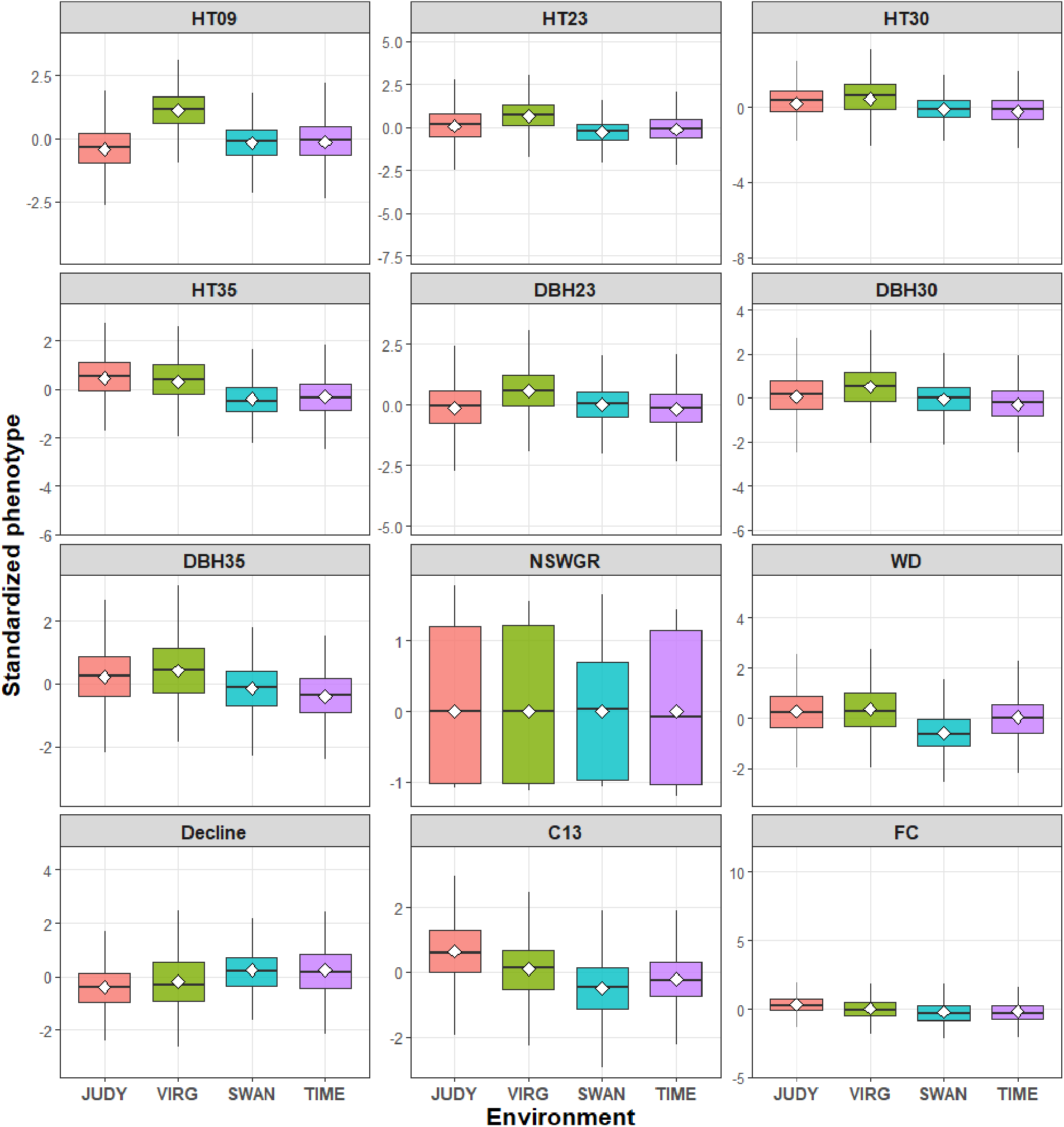
Boxplot of standarized phenotipic data for the 12 traits studied in the lodgepole pine population in four enviroment. See text for trait and enviroment abbrevations.

Integrating multiple omics layers into GP presents an important analytical challenge because these datasets differ substantially in dimensionality, scale, and structure (Picard *et al*., 2021; Montesinos-López *et al*., 2025). Moreover, the relationship within and among genomic, environmental, and other omics layers are often nonlinear and difficult to capture using conventional linear models. Machine learning (ML) and kernel-based approaches provide flexible alternatives for modeling such complex relationships (Yan and Wang, 2023; Crossa *et al*., 2025). While linear kernels form the basis of traditional GBLUP models, nonlinear kernels such as Gaussian and arc-cosine functions can capture more complex patterns, including gene-gene interactions (Alves *et al*., 2019; Cappa *et al*., 2026) and nonlinear G×E interactions (De Los Campos *et al*., 2010; Cuevas *et al*., 2016, 2017; Bandeira e Sousa *et al*., 2017; Crossa *et al*., 2019; Costa-Neto *et al*., 2020).

Recent studies in crops have demonstrated that integrating genomic, metabolomic, transcriptomic, and phenomic information can improve prediction accuracy when combined with advanced ML approaches (Wu *et al*., 2024; Amin *et al*., 2025; Montesinos-López *et al*., 2025; Mohamedikbal *et al*., 2025). However, relatively few studies have evaluated multi-omics prediction in multi-environment settings (Hu *et al*., 2021), despite growing recognition that robust multi-omics frameworks should be validated across diverse environments and incorporate G×E interactions (Montesinos-López *et al*., 2025; Yoshioka *et al*., 2026). Moreover, applications of these approaches in forest tree breeding remain limited (Wang *et al*., 2023). To our knowledge, no previous study has jointly integrated genomic, enviromic, and metabolomic information within multi-environment ssGBLUP reaction norm models using both linear and nonlinear kernels in forest tree breeding. In this study, we conducted a comprehensive evaluation of multi-environment ssGBLUP reaction norm models integrating genomic, enviromic (i.e., climatic and edaphic), and targeted metabolomic information to predict 12 productivity- and climate-adaptation-related traits measured across four open-pollinated lodgepole pine progeny trials. Specifically, our objectives were to: (i) assess the relative contribution of environmental relationship matrices derived from different omics covariates, and their interactions with genotypes, modeled using alternative linear and nonlinear kernels, in explaining phenotypic variation across environments; and (ii) evaluate the predictive ability of these ssGBLUP reaction norm models for prediction in untested (new) environments under variable climatic conditions.

## Material and Methods

### Genetic material and trial description

Genetic material and field trials used in this study were previously described by (Cappa, Chen, *et al*., 2022). Briefly, the genetic material originated from four open-pollinated (OP) lodgepole pine progeny test trials established under Alberta Forestry and Park’s Region C lodgepole pine breeding program (FGRMS, 2016) and currently managed by Blue Ridge Lumber. The study population was comprised of 224 OP families derived from five natural stands (provenances); Deer Mountain, Inverness River, Judy Creek, Swan Hills, and Virginia Hills, all located within the Region C breeding region. These progeny tests were established at four sites in Alberta: Judy Creek (JUDY; 54°26’42.4’’ N, 115°34’10.7’’ W, 1097 m elevation), Virginia Hills (VIRG; 54°28’26.2’’ N, 115°50’42.1’’ W, 1127 m elevation), Swan Hills (SWAN; and 54°42’43.9’’ N, 115°29’54.0 W, 1033 m elevation), and Timeau (TIME; 54°43’49.6’’ N, 115°17’57.6’’ W, 1064 m elevation). All trials followed a “set in replications” experimental design consisting of five replications (blocks), 21 sets per replication, and 4-tree row plots planted at a spacing of 2.5 × 2.5 m. All trial sites were fenced and surrounded by a single buffer row of lodgepole pine seedlings planted around the perimeter to minimize edge effects. Additional information on site characteristics and experimental design is provided in (Cappa, Ratcliffe, *et al*., 2022) (Table 1), while the geographic distribution of the test sites across Alberta, Canada, is illustrated in (Cappa, Chen, *et al*., 2022) (Supplementary Information Fig. S2).

**Table 1.**
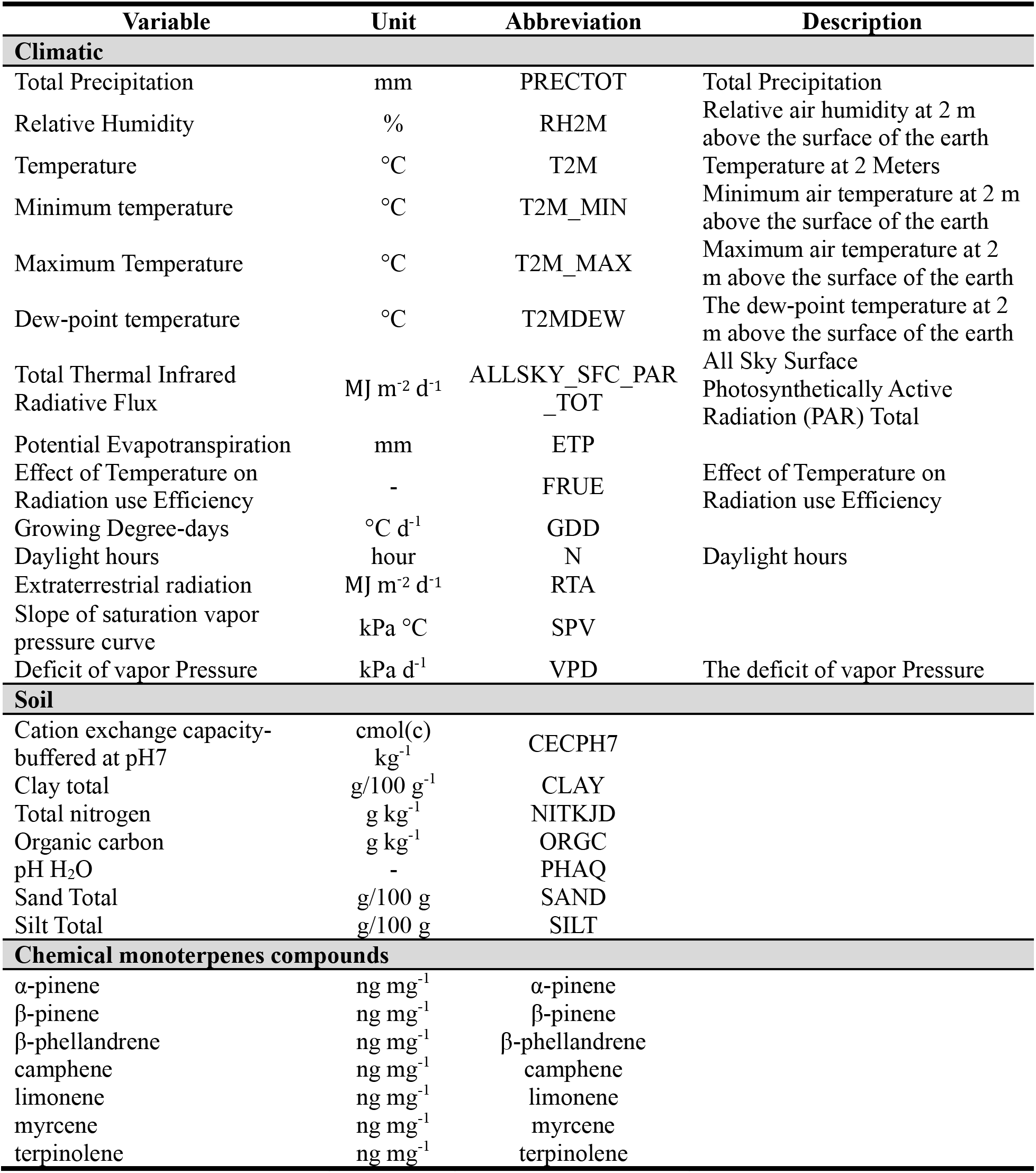
List of covariates considered in this study including climatic, soil and targeted monoterpenes compounds, units, abbreviations and descriptions.

| Variable | Unit | Abbreviation | Description |
|---|---|---|---|
| **Climatic** |  |  |  |
| Total Precipitation | mm | PRECTOT | Total Precipitation |
| Relative Humidity | % | RH2M | Relative air humidity at 2 m above the surface of the earth |
| Temperature | °C | T2M | Temperature at 2 Meters |
| Minimum temperature | °C | T2M_MIN | Minimum air temperature at 2 m above the surface of the earth |
| Maximum Temperature | °C | T2M_MAX | Maximum air temperature at 2 m above the surface of the earth |
| Dew-point temperature | °C | T2MDEW | The dew-point temperature at 2 m above the surface of the earth |
| Total Thermal Infrared Radiative Flux | MJ m<sup>-2</sup> d<sup>-1</sup> | ALLSKY_SFC_PAR_TOT | All Sky Surface Photosynthetically Active Radiation (PAR) Total |
| Potential Evapotranspiration | mm | ETP |  |
| Effect of Temperature on Radiation use Efficiency | - | FRUE | Effect of Temperature on Radiation use Efficiency |
| Growing Degree-days | °C d<sup>-1</sup> | GDD |  |
| Daylight hours | hour | N | Daylight hours |
| Extraterrestrial radiation | MJ m<sup>-2</sup> d<sup>-1</sup> | RTA |  |
| Slope of saturation vapor pressure curve | kPa °C | SPV |  |
| Deficit of vapor Pressure | kPa d<sup>-1</sup> | VPD | The deficit of vapor Pressure |
| **Soil** |  |  |  |
| Cation exchange capacity-buffered at pH7 | cmol(c) kg<sup>-1</sup> | CECPH7 |  |
| Clay total | g/100 g<sup>-1</sup> | CLAY |  |
| Total nitrogen | g kg<sup>-1</sup> | NITKJD |  |
| Organic carbon | g kg<sup>-1</sup> | ORGC |  |
| pH H<sub>2</sub>O | - | PHAQ |  |
| Sand Total | g/100 g | SAND |  |
| Silt Total | g/100 g | SILT |  |
| **Chemical monoterpenes compounds** |  |  |  |
| α-pinene | ng mg<sup>-1</sup> | α-pinene |  |
| β-pinene | ng mg<sup>-1</sup> | β-pinene |  |
| β-phellandrene | ng mg<sup>-1</sup> | β-phellandrene |  |
| camphene | ng mg<sup>-1</sup> | camphene |  |
| limonene | ng mg<sup>-1</sup> | limonene |  |
| myrcene | ng mg<sup>-1</sup> | myrcene |  |
| terpinolene | ng mg<sup>-1</sup> | terpinolene |  |

### Phenotypic data

The four progeny trials were assessed for two growth productivity traits, diameter at breast height (DBH, measured at 1.3 m above ground) and total tree height (HT). DBH were measured at ages 23, 30, and 35, whereas HT was measured at ages 9, 23, 30, and 35. At age 35, DBH and HT were assessed only in trees selected for genomic sampling (see Genomic Data subsection below). Wood density (WD) was assessed at age 35 years using 5-mm increment cores extracted at DBH (for details see (Cappa, Chen, *et al*., 2022)).

Resistance to *Endocronartium harknessii* (J.P. Moore) Y. Hirats (western gall rust, WGR) was evaluated at age 36 using a qualitative seven-class scoring system ranging from the absence of gall symptoms (class 1) to dead trees (class 7). Because classes 3, 5, and 7 occurred infrequently across the trial sites, they were merged with classes 2, 4, and 6, respectively, resulting in a four-class resistance scale. These four WGR classes were subsequently transformed into normal score (NSWGR) following the method described by (Gianola and Norton, 1981).

The dendrochronological growth decline index (Decline) was derived from tree-ring measurements obtained from the same increment cores used for wood quality assessment. This index quantifies long-term growth reduction associated with repeated drought events by comparing the mean basal area increment (BAI) during a pre-drought reference period (1997-2001) with the mean BAI during the final five years of the study period (2012-2016). Larger decline values indicate greater growth reductions. Further details regarding the calculation and interpretation of this index are provided in (Sebastian-Azcona *et al*., 2025).

The two outer slabs remaining after pneumatic processing of the increment cores for wood density determination, were subsequently used to characterize lifetime variation in stable carbon isotope composition (C13). In addition, the carbon fraction (FC), defined as the proportion of carbon relative to total dry biomass, was measured during elemental analysis. A detailed description of sample preparation and analytical procedures is provided by (Cappa, Klutsch, *et al*., 2022).

Table S1 summarizes all evaluated traits, including the number of observations and descriptive statistics for each trait on their original scale (i.e., prior to standardization). The distribution of the twelve traits across the four environments is illustrated by Figure 1 using boxplots.

### Genomic data

As described by (Cappa, Ratcliffe, *et al*., 2022), 40 OP families were selected from the original population of 224 families, to represent the full range of height variation at age 30, including families with high, intermediate, and low growth performance. Approximately 10 trees per OP family were sampled at each of the four sites, yielding a total of 1,600 sampled individuals (see (Cappa, Ratcliffe, *et al*., 2022) for details). Additionally, 35 candidate forward-selected trees, previously identified based on their height breeding values (BVs) at age 30, and representing an additional 28 OP families, were included in the sequencing efforts. Consequently, a total of 1,635 trees from 68 OP families were selected and needles sampled for genotyping.

Of the 1,635 sampled trees, 1,554 DNA samples passed quality-control procedures after extraction and were subsequently genotyped using the genotyping-by-sequencing (GBS) platform (Chen *et al*., 2013) following the protocol described by (Cappa, Ratcliffe, *et al*., 2022). After filtering for a minor allele frequency (MAF) ≥ 1% and allowing up to 30% missing genotype data per marker, the final dataset was comprised of 1,490 trees genotyped at 25,099 (25K) single nucleotide polymorphisms (SNPs). Missing genotypes were imputed using the mean allele frequency method described by (Ratcliffe *et al*., 2015). Because a reference genome assembly was unavailable for lodgepole pine at the time oof analysis, SNP calling and genotyping were conducted using the reference-free UNEAK pipeline.

### Enviromic data

Climatic environmental covariates were obtained using the get_weather() function from the R package envRtype (Costa-Neto *et al*., 2021). Fourteen climatic variables (Table 1) were retrieved from NASA POWER satellite-derived datasets using the geographic coordinates of each trial site. Climate records covering the period from plantation establishment (between May 31 and June 18, 1982) (see Table 1 in (Cappa, Ratcliffe, *et al*., 2022)) to the final measurement dates (July 12 to 20, 2017). To better represent conditions influencing tree growth, only data corresponding to the active growing season (April 15 to October 15) were retained (Chhin *et al*., 2008). The resulting 35-year climate dataset was used to characterize historical environmental variability across sites and to approximate the range of climate conditions that trees may encounter in future deployment environments (Gillberg *et al*., 2019).

Soil environmental covariates were obtained for each site using the get_soil() function from the R package SoilType (Fritsche-Neto, 2024). This package also uses the geographic coordinates of the trial sites to retrieve soil information from global databases, particularly the World Soil Information Service (WoSIS). Based on the WoSIS, seven physical and chemical soil covariates (Table 1) were obtained for the locations closest to each experimental site.

### Targeted metabolomic data

Secondary metabolites, primarily monoterpenes associated with tree defense, were identified and quantified from phloem tissue collected between May and July 2017 across the four lodgepole pine test sites. A detailed description of the sampling, extraction, and analytical procedures are described in (Cappa, Chen, *et al*., 2022). In total, 15 hexane-extractable monoterpenes were identified. From these, seven compounds (Table 1) were selected for further analysis, based on their relatively high abundance in the phloem and their documented role in lodgepole pine defense against *Dendroctonus ponderosae* Hopkins (mountain pine beetle) (Erbilgin, 2019). These compounds were used as targeted monoterpene covariates in subsequent analyses.

Although monoterpenes are metabolomic traits, they were incorporated into the reaction norm framework as biologically informed descriptors of environmental similarity. As outlined in the Introduction, metabolite profiles reflect the combined influence of genetic and environmental conditions. Consequently, targeted monoterpenes may capture adaptive and stress-related processes that are not fully represented by conventional climatic and soil variables. Thus, these metabolite profiles provide offering complementary information for characterizing environmental heterogeneity and genotype-by-environment interactions.

### Quantitative genetics analyses

#### Pedigree-based multi-environment mixed model

To obtain heritability estimates and additive genetic correlations across sites, a classical pedigree-based multi-environment mixed model (ABLUP) was fitted for each trait as follows:

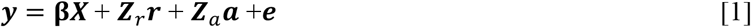

where, ***y*** is the vector of individual tree phenotypes; **β** is the vector of fixed effects of genetic group nested within site formed according to the five provenances; ***r*** is the vector of random replicate effect nested within site; ***a*** is the random vector of additive genetic effects (*i.e.*, breeding values) distributed as ***a***∼***N***(**0**, ***Σ_a_***⊗***A***), where ***Σ_a_*** is the multi-environment genetic effects (co)variance matrix and ***A***-matrix is the average numerator relationship matrix derived from the pedigree information (Henderson, 1984). Finally, ***e*** is the vector of random residual distributed as ***e***∼***N***(**0**, ***R*_0_**⊗***I***), where ***R*_0_** is the residual (co)variance matrix for the four environments (sites) with dimension 4 × 4, and ***I*** denotes the identity matrix, while the symbol ⊗ stands for the Kronecker operator. We assumed an unstructured (co)variance matrix for the genetic effects (***Σ_a_***). However, given that the sites were assessed separately, the residual covariance across sites is assumed to be zero. The matrices ***X***, ***Z****_r_*, and ***Z****_a_* relate the observation to the means of the site effects in **β**, the replicate effects in ***r***, and the additive genetic effects for each tree in ***a***.

Individual narrow-sense heritability estimates for each environment *i* 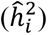 and genetic correlations between environment *i* and *j* (*r̂_ai_*_j_), were estimated as follows:

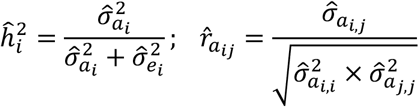

where, 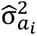 is the estimated variance for the additive genetic effects, and 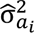 is the estimated residual error for environment *i*. *σ*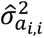 and 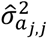 are the estimated additive genetic variances in the environment *i* and *j*, respectively, and 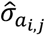 is the estimated additive genetic covariance between environments *i* and *j*.

Variance components, narrow-sense heritability estimates and genetic correlations between sites from model [1] (and their standard errors), were estimated using average information restricted maximum likelihood (AI-REML) with the ASReml-R package (version 4.2.0) (Butler *et al*., 2023).

#### Genomic-pedigree-based multi-environment ssGBLUP predictive mixed models

To assess the impact of multi-omic covariates and their interactions with the environment on the proportion of variance explaining and genomic prediction, five hierarchical ssGBLUP prediction models were fitted following the (Jarquín *et al*., 2014) approach.

Assuming that phenotypic data were collected from *q* trees evaluated across *r* sites. Trees may have different means depending on their membership in different genetic groups, and replicates were nested within sites. Let *i* (*i* = 1, 2, …, *r*) denote the subscript for site or environment, *j* (*j* = 1, 2, …, *q*) the subscript for tree, *k* (*k* = 1, 2, …, *g*) the corresponding genetic group, and *l* (*l* = 1, 2, …, *s*) the replicate within site. Let *y_i_*_j*l*_ and *e_i_*_j*l*_ denote the observed phenotype and the residual effect of individual *j* evaluated at site *i*, and replicate *l*, respectively. The first ssGBLUP model included a genetic group (β*_k_*) as fixed effects, together with random environment effects (*E_i_*), random replicate within site effects (*R_l_*_(*i*)_), and additive genetic value (*a*_j_). Then, the first ssGBLUP mixed model is defined as:

Model 1 (M1, E + G)

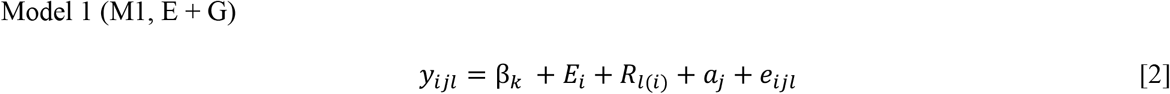

where the assumed distributions are: 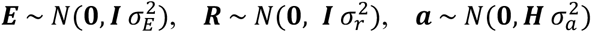 and 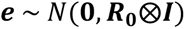. The variance components 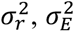 and 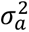, represent the variances associated with each effect, assumed to be common across environments. Matrices ***R*_0_** and ***I*** were defined previously, and ***H*** corresponds to the combined pedigree-genomic relationship matrix. The inverse of the relationship matrix that combines combined pedigree- and marker information (***H*^−1^**) was derived by (Legarra *et al*., 2009; Aguilar *et al*., 2010; Christensen *et al*., 2012), and calculated for each ***G***-matrix as:

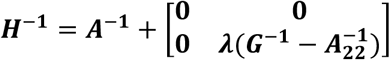

where ***λ*** scales the differences between genomic and pedigree-based information, ***G*^−1^** is the inverse of the genomic-based relationship matrices, and 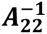 is the inverse of the pedigree-based relationship matrix for the genotyped individuals only (the subscript 2 denotes genotyped individuals). The weighting factor λ was set to 0.95 for all ssGBLUP models. The ***G***-matrix was estimated following the first method proposed by (VanRaden, 2008):

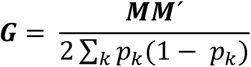

where, ***M*** is the centered matrix of SNP covariates, and *p_k_* is the current (or observed) allele frequency of the genotyped trees for marker *k*. The ***G***-matrix was scaled to have the same diagonal and off-diagonal averages as the corresponding ***A****_22_*-matrix, according to (Christensen *et al*., 2012) (eq. 4). Finally, in the model [2], the fixed-effect vector **β** represents the systematic effects associated with the *k* genetic groups.

Model 2 (M2, E + W + G)

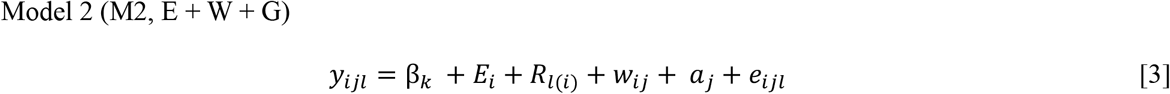

Model M2 extends Model M1 (Equation [2]) by incorporating a new random effect associated with environmental covariates, including climatic, soil, or targeted monoterpene covariables (***w***). This effect is assumed to follow 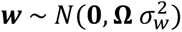, where **Ω** represents the environmental kernel, i.e., the relationship matrix among environments (sites) derived from climatic, soil, or monoterpene information using linear and nonlinear kernel functions (see below), and 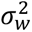 is the variance associated with the environmental covariates.

Model 3 (M3, E + G + W + G×W)

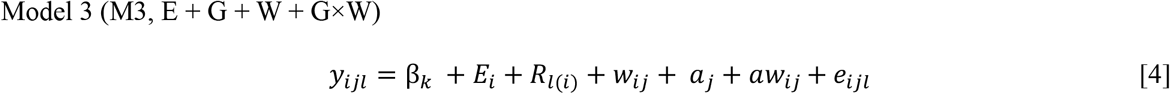

Model M3 incorporates the random interaction vector between additive genetic effects (***a***) and environmental covariates (***w***). This effect is modeled as 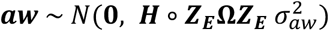, where 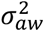 is the variance associated with the genotype × environmental covariate interaction, ***Z_E_*** is the incidence matrix for site (or environment) effects, and ∘ denotes the Hadamard product of matrices.

Model 4 (M4, E + G + W + G×E)

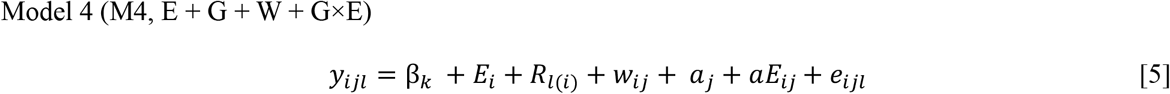

Model M4 incorporates the random interaction effect between additive genetic effects (***a***) and the evaluation sites 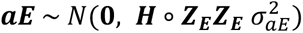, where 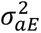 is the variance associated with the genotype × enviroment (G×E) interaction.

Model 5 (M5, E + G + W + G×E + G×W)

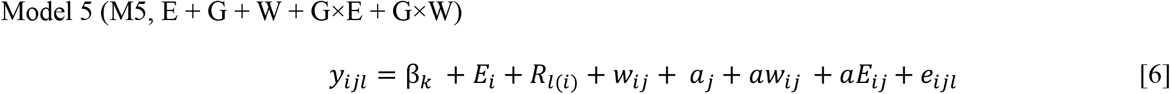

Finally, Model M5 simultaneously includes both interaction effects: genotype × environment (G×E) (Equation [5]) and genotype × environmental covariates (G×W) (Equation [4]).

Table 2 summarizes the five multi-environment ssGBLUP models evaluated across the 12 traits, including the covariates, kernel types, model abbreviations, and corresponding model formulations. In total, 37 model-covariate-kernel combinations were assessed.

**Table 2.** Summary of the multi-environment ssGBLUP models evaluated for each of the twelve traits, including order, covariates, kernel type, model-kernel-covariate abbreviations, and the corresponding model. Kernels, and covariates are described in the text. Model abbreviations include the following components: environment effects (S), additive genetic effects (G), environment covariates effects (W), genotype-by-environment interaction (G×E), and genotype-by-covariate interaction (G×W). Covariates correspond to climatic (c), soil (s), and chemical monoterpene (m) data. Nonlinear kernels include the Gaussian kernel (GK) and deep arc-cosine kernel (AK) approaches. While the deep arc-cosine kernel (AK) was fitted using a single layer (1), the Gaussian kernel (GK) was fitted using a bandwidth parameter equal to 1 (*h* = 1).

| Order | Covariates | Kernel | Abbreviation | Model |
|---|---|---|---|---|
| 1 | - | - | M1 | E + G |
| 2 | Climatic | Linear | M2 | E + G + Wc |
| 3 | Climatic | Nonlinear GK | M2 | E + G + Wc (GK) |
| 4 | Climatic | Nonlinear AK | M2 | E + G + Wc (AK) |
| 5 | Soil | Linear | M2 | E + G + Ws |
| 6 | Soil | Nonlinear GK | M2 | E + G + Ws (GK) |
| 7 | Soil | Nonlinear AK | M2 | E + G + Ws (AK) |
| 8 | Monoterpenes | Linear | M2 | E + G + Wm |
| 9 | Monoterpenes | Nonlinear GK | M2 | E + G + Wm (GK) |
| 10 | Monoterpenes | Nonlinear AK | M2 | E + G + Wm (AK) |
| 11 | Climatic | Linear | M3 | E + G + Wc + G×Wc |
| 12 | Climatic | Nonlinear GK | M3 | E + G + Wc + G×Wc (GK) |
| 13 | Climatic | Nonlinear AK | M3 | E + G + Wc + G×Wc (AK) |
| 14 | Soil | Linear | M3 | E + G + Ws + G×Ws |
| 15 | Soil | Nonlinear GK | M3 | E + G + Ws + G×Ws (GK) |
| 16 | Soil | Nonlinear AK | M3 | E + G + Ws + G×Ws (AK) |
| 17 | Monoterpenes | Linear | M3 | E + G + Wm + G×Wm |
| 18 | Monoterpenes | Nonlinear GK | M3 | E + G + Wm + G×Wm (GK) |
| 19 | Monoterpenes | Nonlinear AK | M3 | E + G + Wm + G×Wm (AK) |
| 20 | Climatic | Linear | M4 | E + G + Wc + G×E |
| 21 | Climatic | Nonlinear GK | M4 | E + G + Wc + G×E (GK) |
| 22 | Climatic | Nonlinear AK | M4 | E + G + Wc + G×E (AK) |
| 23 | Soil | Linear | M4 | E + G + Ws + G×E |
| 24 | Soil | Nonlinear GK | M4 | E + G + Ws + G×E (GK) |
| 25 | Soil | Nonlinear AK | M4 | E + G + Ws + G×E (AK) |
| 26 | Monoterpenes | Linear | M4 | E + G + Wm + G×E |
| 27 | Monoterpenes | Nonlinear GK | M4 | E + G + Wm + G×E (GK) |
| 28 | Monoterpenes | Nonlinear AK | M4 | E + G + Wm + G×E (AK) |
| 29 | Climatic | Linear | M5 | E + G + Wc + G×E + G×Wc |
| 30 | Climatic | Nonlinear GK | M5 | E + G + Wc + G×E + G×Wc (GK) |
| 31 | Climatic | Nonlinear AK | M5 | E + G + Wc + G×E + G×Wc (AK) |
| 32 | Soil | Linear | M5 | E + G + Ws + G×E + G×Ws |
| 33 | Soil | Nonlinear GK | M5 | E + G + Ws + G×E + G×Ws (GK) |
| 34 | Soil | Nonlinear AK | M5 | E + G + Ws + G×E + G×Ws (AK) |
| 35 | Monoterpenes | Linear | M5 | E + G + Wm + G×E + G×Wm |
| 36 | Monoterpenes | Nonlinear GK | M5 | E + G + Wm + G×E + G×Wm (GK) |
| 37 | Monoterpenes | Nonlinear AK | M5 | E + G + Wm + G×E + G×Wm (AK) |

#### Linear and nonlinear environmental and targeted metabolomic kernels

Climatic and soil environmental data, together with targeted metabolomic covariates (monoterpenes), were used to estimate the environmental similarity matrix among environments (**Ω**). In this study, three methods were evaluated to estimate environmental similarity kernels. As a reference, a linear kernel was computed from the environmental covariate matrix ***W***, which included climatic, soil, and monoterpene variables. In this case, ***W*** corresponds to a centered and standardized matrix of covariates with dimensions equal to the number of sites (*r*) by the number of covariates (*m*) (*m* = 14 for climatic variables and *m* = 7 for soil and targeted monoterpenes) (Table 1). The linear relationship matrix (Linear) was then calculated as:

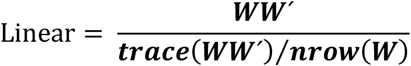

Additionally, two nonlinear kernels from machine learning were evaluated to capture more complex relationships among environments: the Gaussian kernel (GK) (Gianola *et al*., 2006; Gianola and Van Kaam, 2008) and the arc-cosine kernel (AK) (Cuevas *et al*., 2019).

The GK estimates similarity between environments based on the Euclidean distance between their covariate vectors, allowing the model to account for nonlinear patterns that cannot be captured by linear kernels. A bandwidth parameter (*h*) controls the rate at which similarity decays with increasing distance between environments. In this study, four different *h* values (0.1, 0.25, 0.5, and 1) were initially evaluated, and the final analyses were conducted using *h* = 1.

The Linear and GK relationship kernels were calculated using the W_matrix() function from the R package envRtype (Costa-Neto *et al*., 2021).

The AK was also evaluated following the approach proposed by (Cuevas *et al*., 2019), originally introduced by (Cho and Saul, 2009). This kernel measures similarity between environments based on the angle between their environmental covariate vectors, allowing the model to capture complex nonlinear and hierarchical relationships. The arc-cosine kernel can be interpreted as an approximation to computations performed by deep neural networks, enabling flexible representations of interactions among environmental variables. In this study, the AK was computed from the environmental covariate matrix ***W*** using a single layer (layer = 1), as implemented by (Cuevas *et al*., 2019).

Following to (Prado *et al*., 2024), the environmental similarity structures (**Ω**) generated by the linear and nonlinear kernels were compared with the pattern of the pedigree-based genetic correlations among sites for each trait. For each covariate source (climatic, soil, and targeted monoterpene data), pairwise environmental similarities were calculated, and their distributions were visualized using density plots.

#### Assessment of the multi-environment ssGBLUP models

First, models M1-M5 (Table 2) were fitted using the complete dataset available for each trait (Table S1). This analysis enabled the estimation of variance components (proportion of variance) and the calculation of goodness-of-fit statistics for model comparison, including the log-likelihood, and Akaike Information Criterion (AIC).

Second, to evaluate the predictive ability of the five proposed ssGBLUP models, a Leave-One-Environment-Out cross-validation scheme was implemented. The objective was to predict the performance of genotypes already observed in a novel (unobserved) climate-variable environment. To mimic that situation, each environment (i.e., site) was treated in turn as a future environment. At each iteration, the training population included phenotypic observations from three environments, which were used to predict the performance of genotypes in the excluded environment (validation). This procedure was repeated across all four environments.

Predictive ability was quantified as the Pearson correlation (*r*) between the predicted genetic values and the corresponding observed values in the omitted environment.

## Results

### Pedigree-based heritability estimates and additive genetic correlations across sites

Pedigree-based heritability estimates varied across traits and sites (Table 3). Growth traits showed a clear age-dependent pattern, with moderate heritability estimates at early ages (e.g., HT09: 0.19-0.38; DBH23: 0.18-0.29) that increased substantially at later ages (e.g., HT35: 0.72-0.87; DBH35: 0.33-0.61). Among traits, HT35, C13, and WD exhibited the highest average heritability estimates (0.80, 0.59, and 0.46, respectively), whereas DBH at younger ages and Decline showed lower values, with Decline presenting particularly low and variable estimates across sites (0.00-0.41). Across environments, SWAN and JUDY tended to show slightly higher heritability estimates (0.40 and 0.42, respectively) on average compared to VIRG and TIME. It should be noted that HT35 and DBH35 were assessed only in the genomic sampling subset; therefore, the relatively high heritability estimates observed for these traits should be interpreted in light of this sampling design.

**Table 3.** Estimated heritabilities (with their approximate standard error) for the 12 traits evaluated in the lodgepole population using a pedigree-based relationship matrix (*A*-matrix). Abbreviations used for the traits and sites are described in the text.

| Traits/sites | JUDY | VIRG | SWAN | TIME | Average |
|---|---|---|---|---|---|
| **HT09** | 0.30 (0.06) | 0.27 (0.07) | 0.38 (0.05) | 0.19 (0.04) | **0.29 (0.06)** |
| **HT23** | 0.42 (0.07) | 0.28 (0.07) | 0.46 (0.06) | 0.35 (0.05) | **0.38 (0.06)** |
| **HT30** | 0.35 (0.06) | 0.33 (0.07) | 0.42 (0.06) | 0.33 (0.05) | **0.36 (0.06)** |
| **HT35** | 0.72 (0.21) | 0.85 (0.25) | 0.87 (0.21) | 0.76 (0.21) | **0.80 (0.22)** |
| **DBH23** | 0.18 (0.05) | 0.29 (0.07) | 0.23 (0.04) | 0.18 (0.04) | **0.22 (0.05)** |
| **DBH30** | 0.22 (0.06) | 0.30 (0.07) | 0.25 (0.04) | 0.20 (0.04) | **0.24 (0.05)** |
| **DBH35** | 0.61 (0.21) | 0.57 (0.22) | 0.33 (0.16) | 0.33 (0.16) | **0.46 (0.19)** |
| **NSWGR** | 0.43 (0.07) | 0.51 (0.08) | 0.55 (0.06) | 0.49 (0.06) | **0.50 (0.07)** |
| **WD** | 0.65 (0.22) | 0.38 (0.21) | 0.33 (0.16) | 0.48 (0.19) | **0.46 (0.20)** |
| **Decline** | 0.41 (0.20) | 0.00 (0.00) | 0.24 (0.16) | 0.23 (0.17) | **0.22 (0.13)** |
| **C13** | 0.65 (0.20) | 0.53 (0.19) | 0.63 (0.19) | 0.54 (0.17) | **0.59 (0.19)** |
| **FC** | 0.58 (0.21) | 0.24 (0.18) | 0.22 (0.14) | 0.17 (0.14) | **0.30 (0.17)** |
| **Average** | **0.42 (0.14)** | **0.34 (0.11)** | **0.40 (0.12)** | **0.33 (0.12)** | **0.37 (0.12)** |

Genetic correlations among sites were positive but varied considerably depending on the trait (Figure 2). Early-age growth traits showed the lowest and most variable correlations across environments (HT09: 0.25-0.71, mean = 0.50; HT23: 0.55-0.70, mean = 0.65; HT30: 0.50-0.75, mean = 0.56), indicating a stronger genotype-by-environment (G × E) interaction at younger ages. Similarly, Decline exhibited relatively low, and highly variable, correlations (0.19-1.00; mean = 0.52), further supporting substantial G × E effects for this trait. In contrast, later-age traits such as HT35 (0.70-1.00; mean = 0.90) and DBH35 (0.52-1.00; mean = 0.81) showed higher and more consistent correlations across sites, indicating greater stability of genetic effects over time. However, both traits were assessed only in the genomic sampling subset at age 35. Intermediate patterns were observed for DBH23 (0.49-0.83; mean = 0.70) and wood density (WD: 0.26-1.00; mean = 0.76), suggesting more moderate levels of G × E interaction. Notably, lower correlations were more frequently observed between TIME and JUDY or VIRG (the more distant sites) (e.g., HT09: 0.25 and 0.34; Decline: 0.19 and 0.44), whereas correlations between TIME and SWAN were generally higher (e.g., HT09: 0.71; Decline: 0.86).

**Figure 2.**
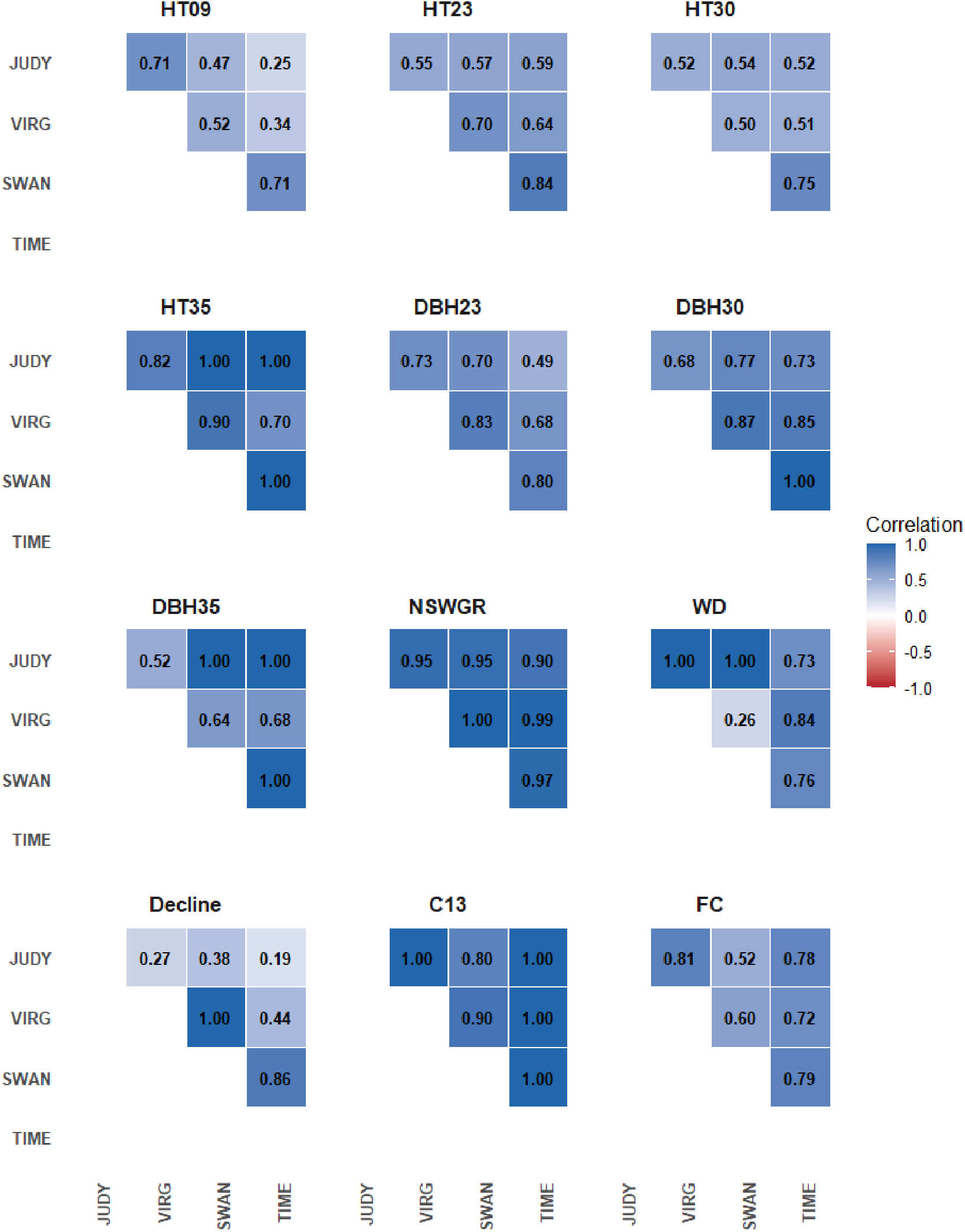
Estimated genetic correlations between sites for each trait from the multiple-site model using pedigree-(*A*-matrix) relationship matrices for the lodgepole pine population. The dark red to dark blue color of each individual cell above the diagonal reflects the strength of the genetic correlation. Abbreviations used for the traits and sites are described in the text.

### Covariate patterns across sites

The ***W*** matrices show differences in the clustering patterns of environments across the three sets of covariates (Figure 3). For climatic variables (A), TIME was separated from the other sites, showing contrasting patterns for most temperature-and precipitation-related covariates, while SWAN, VIRG, and JUDY clustered more closely, indicating more similar climatic conditions among them. In contrast, soil covariates (B) showed a different clustering pattern, where VIRG and JUDY grouped together, suggesting similar soil composition, whereas SWAN and TIME appeared more differentiated from both each other and the VIRG–JUDY group. For targeted monoterpenes (C), TIME again showed a markedly different profile, with consistently similar values across all compounds, separating from the remaining sites. Overall, these results highlight that environmental similarity among sites depends on the type of covariates considered, with TIME consistently representing the most distinct environment.

**Figure 3.**
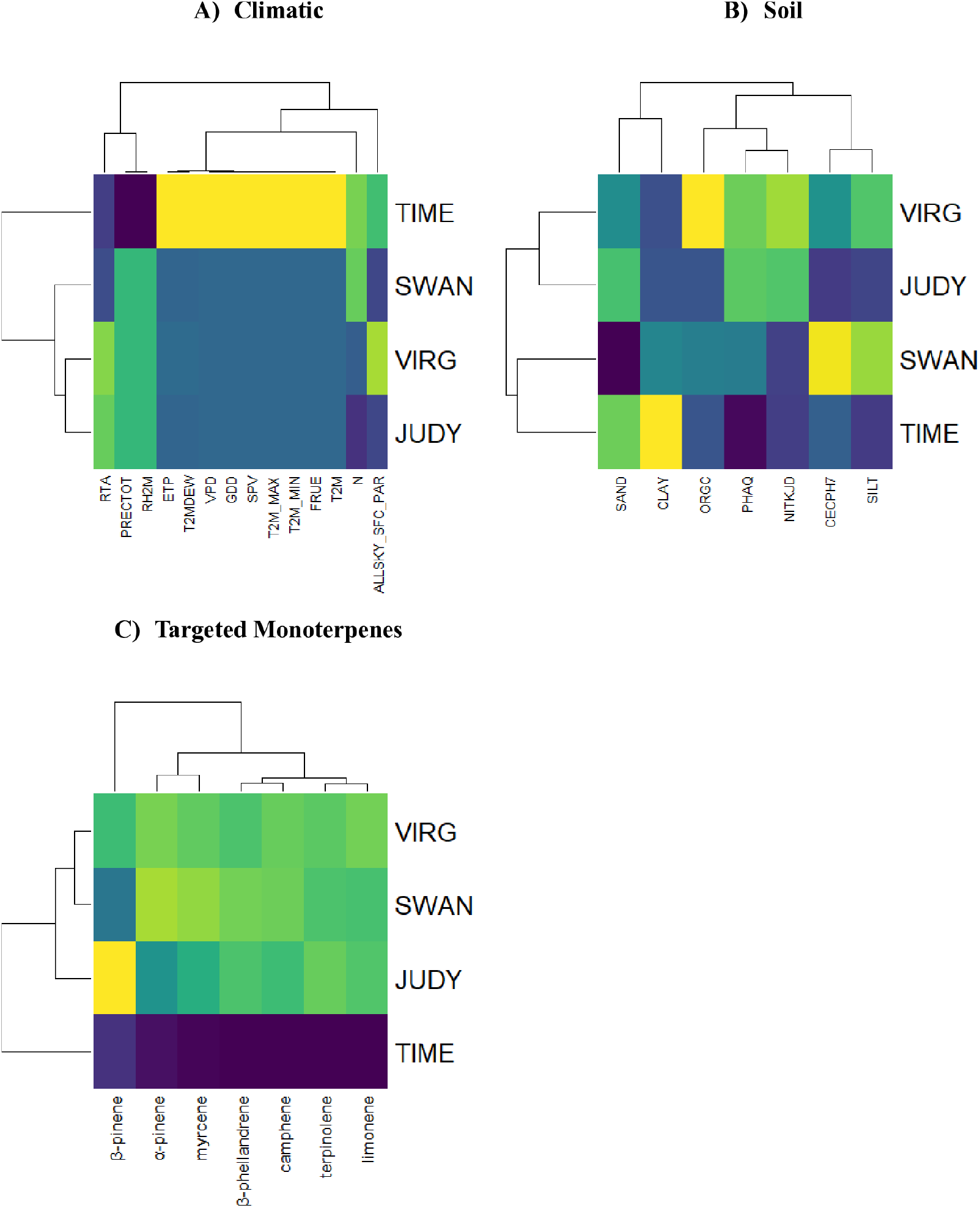
Heta maps of the climatic (A), soil (B) and targeted monoterpenes (C) covariates across the four lodgepole pine environments. Covariate abbreviations are provided in Table 1, and site abbreviations are described in the text.

### Environmental similarity patterns across covariates and kernel methods

The environmental similarity patterns across sites varied depending on both the type of covariates and the kernel used (Figure 4). For climatic covariates (A), the linear kernel showed moderate similarities among SWAN, VIRG, and JUDY, with TIME remaining clearly distinct, a pattern that was further accentuated under the nonlinear kernels (GK and AK), which increased the contrast among environments. For soil covariates (B), the linear kernel suggested closer similarity between VIRG and JUDY, while SWAN and TIME were more differentiated; this structure was partially preserved under GK, although similarities became more polarized, whereas AK highlighted sharper contrasts across all sites. In the case of targeted monoterpenes (C), TIME consistently appeared as the most dissimilar environment across all kernels, although the strength of the pairwise dissimilarities differed among kernels. In summary, the nonlinear kernels produced clustering patterns that differed from those obtained with the linear kernel and, in some cases, highlighted environmental contrasts more clearly. Nevertheless, the overall relationships among sites were broadly consistent across kernel approaches.

**Figure 4.**
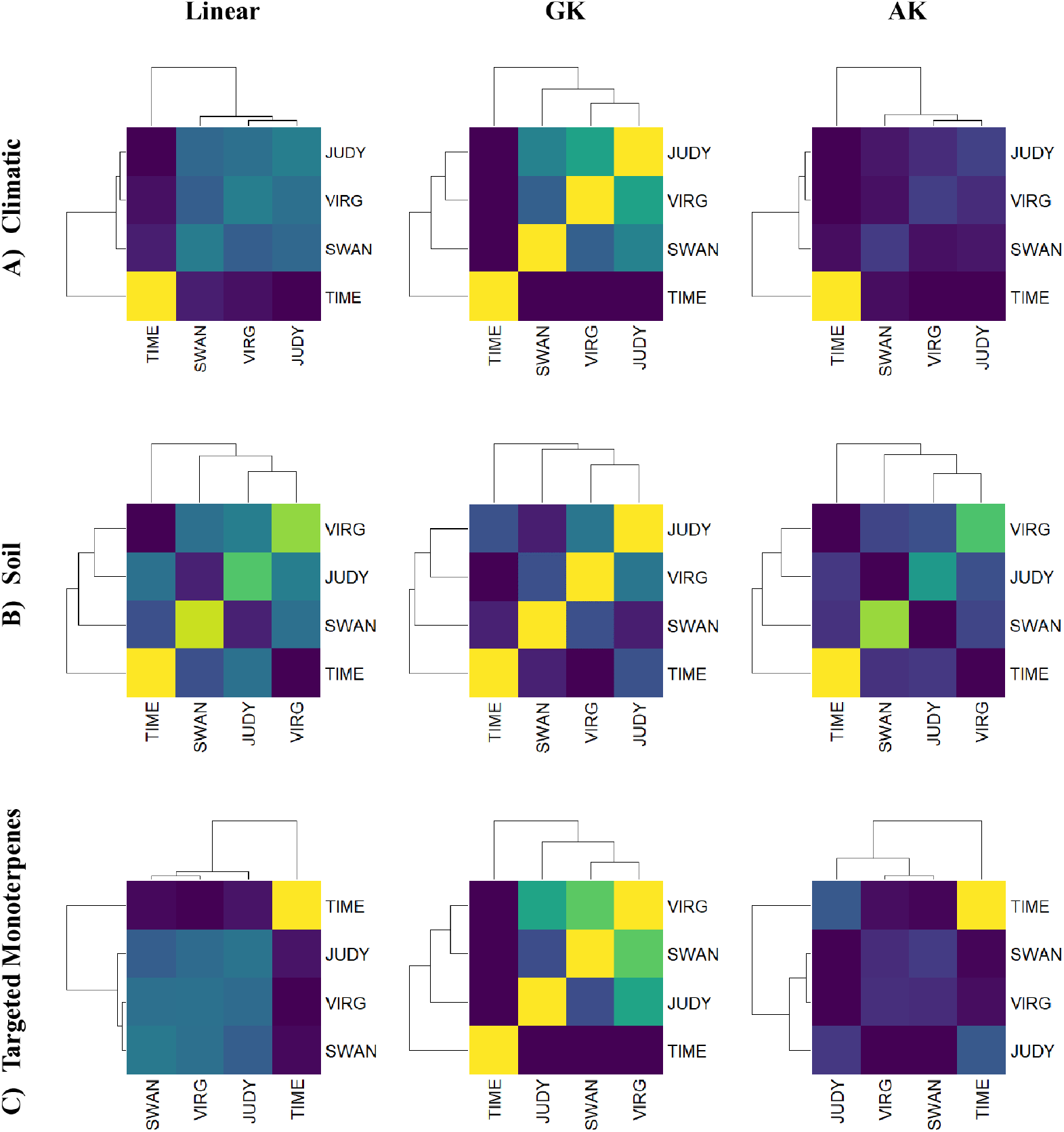
Heat maps of the environment similarity matrix among the four lodgepole pine environments considering climatic (A), soil (B) and targeted monoterpenes (C) covariates, estimated using a linear kernel (Linear) and two nonlinear kernels: the Gaussian kernel (GK; bandwidth parameter *h* = 1) and the deep arc-cosine kernel (AK; one layer). Site abbreviations are described in the text.

The environmental kernels derived from climatic, soil, and targeted monoterpene covariates generally displayed broader similarity distributions than the G×E interaction estimated from the pedigree-based additive genetic correlations (Figure S1). This indicates that environmental covariates capture a wider spectrum of environmental heterogeneity among sites, whereas genetic correlations tend to be concentrated within a narrower range of similarities. Consequently, the G×E structure inferred from the genetic correlations may only partially reflect the environmental contrasts present across these multi-environment lodgepole pine trials, potentially masking part of the underlying environmental heterogeneity.

### Model fit of multi-environment ssGBLUP models across traits

Complete log-likelihood (logL) and Akaike Information Criterion (AIC) values for all evaluated traits, models, kernel functions, and covariate types are presented in Table S2. Overall, the effect of including interaction terms on model fit was trait-dependent (Table 4). Models incorporating interactions yielded lower AIC values for several traits, particularly early growth and Decline-related traits, whereas the baseline model (M1: E + G) remained competitive or achieved the lowest AIC for others. In particular, the inclusion of G×E interaction (M4) produced the largest reduction in AIC for early growth traits (HT09, HT23, HT30, DBH23) and for Decline. Notably, these traits also exhibited lower to moderate genetic correlations among sites (Figure 2), consistent with a stronger influence of a G×E interaction. In contrast, traits with higher and more stable genetic correlations across sites (e.g., HT35 and DBH35) showed little to no improvement, with negligible AIC differences among models. The addition of genotype-by-environmental covariate interaction (G×W; M3) led to moderate improvements in some traits, but the full model (M5) did not consistently outperform M4, indicating limited additional improvement in model fit from including both interaction terms simultaneously. For traits with stronger G×E effects (HT09, HT23, HT30), M5 showed AIC values very similar to M4, although slightly higher, indicating no additional improvement over M4.

**Table 4.** Model fit statistics for all evaluated multi-environment ssGBLUP models across traits, including log-likelihood (LogL) and Akaike Information Criterion (AIC). Values are reported for each trait and model, with both LogL and AIC representing averages across kernel types and covariate sets. Lower AIC (highlighted in bold) indicating better model fit. Trait and model abbreviations are described in the main text.

|  | **HT09** | **HT23** | **HT30** | **HT35** | **DBH23** | **DBH30** | **DBH35** | **NSWGR** | **WD** | **Decline** | **C13** | **FC** |
|---|---|---|---|---|---|---|---|---|---|---|---|---|
| **LogL** |  |  |  |  |  |  |  |  |  |  |  |  |
| **M1:** E+G | -3513.9 | -4389.4 | -4681.9 | -517.7 | -4971.8 | -4950.6 | -635.2 | -5055.3 | -585.6 | -586.2 | -525.9 | -690.1 |
| **M2:** E+G+W | -3513.8 | -4389.4 | -4681.9 | -517.7 | -4971.8 | -4950.6 | -635.2 | -5055.3 | -585.0 | -586.2 | -525.9 | -690.1 |
| **M3**: E+G+W+G×W | -3499.1 | -4382.1 | -4669.2 | -517.7 | -4969.8 | -4950.2 | -635.2 | -5054.9 | -584.4 | -585.2 | -525.9 | -689.5 |
| **M4:** E+G+W+G×E | -3487.7 | -4375.2 | -4659.7 | -517.7 | -4967.3 | -4949.3 | -635.2 | -5053.8 | -584.1 | -583.8 | -525.9 | -688.7 |
| **M5:** E+G+W+G×E+G×W | -3487.7 | -4375.2 | -4659.7 | -517.7 | -4967.3 | -4949.3 | -635.2 | -5053.8 | -584.0 | -583.8 | -525.9 | -688.7 |
| **AIC** |  |  |  |  |  |  |  |  |  |  |  |  |
| **M1:** E+G | 7041.8 | 8792.8 | 9377.8 | **1049.4** | 9957.6 | **9915.2** | 1284.3 | **10124.6** | **1185.2** | 1186.5 | **1065.9** | **1394.2** |
| **M2:** E+G+W | 7043.6 | 8794.8 | 9379.8 | 1051.4 | 9959.6 | 9917.2 | 1286.3 | 10126.6 | 1186.1 | 1188.5 | 1067.8 | 1396.2 |
| **M3**: E+G+W+G×W | 7016.2 | 8782.3 | 9356.5 | 1053.4 | 9957.6 | 9918.5 | 1288.3 | 10127.8 | 1186.9 | 1188.4 | 1069.8 | 1397.0 |
| **M4:** E+G+W+G×E | **6993.4** | **8768.4** | **9337.3** | 1053.4 | **9952.6** | 9916.5 | **1288.3** | 10125.7 | 1186.1 | **1185.6** | 1069.8 | 1395.4 |
| **M5:** E+G+W+G×E+G×W | 6995.4 | 8770.4 | 9339.3 | 1055.4 | 9954.6 | 9918.5 | 1290.3 | 10127.7 | 1188.0 | 1187.6 | 1071.8 | 1397.4 |

### Estimates of variance components across the multi-environment ssGBLUP models

#### Impact of covariates and kernels on the variance attributable to environment

Under the M1 (E + G) model, the main environmental (site) effect accounted for 0.00% to 16.59% of the phenotypic variance (Table S2). The largest site effects were observed for HT09 (16.59%), C13 (9.48%), and HT35 (6.97%), whereas no detectable site effect was found for NSWGR.

The inclusion of covariates (W; M2: E + G + W) led to a moderate but consistent reduction in the magnitude of the main site effect compared with the baseline M1 (E + G) model. Across traits, the average site effect decreased from 5.00% under M1 to 3.97% under M2 (-1.03 percentage points (20.48%)), indicating that part of the environmental variance previously attributed to site was captured by explicit covariates. On average, site effects were lower when soil (3.40%) or climate (4.05%) covariates were included than when monoterpene covariates were fitted (4.48%).

The impact of W on the variance attributable to environment was strongly trait-dependent. For instance, in HT09 the site effect dropped from 15.32% (M1) to 0.00% when soil covariates were included, suggesting a near-complete absorption of site differences by soil descriptors. A similar pattern was observed for DBH23, where the site effect was also reduced to 0.00% under soil covariates using nonlinear kernels. In contrast, for traits such as HT30 and DBH30, site variance remained virtually unchanged across all covariates and kernels, indicating that the evaluated covariates had limited influence in explaining site effects. For WD, climate covariates led to a complete reduction of the site effect (0.00%), whereas soil covariates retained values similar to M1, highlighting that the relevance of environmental descriptors depends on both the trait and the covariate considered.

When further examining site effects, differences among kernels (linear, GK, and AK) were generally modest compared with the impact of including W, although some patterns emerged (Table S2). The linear kernel tended to produce results similar to M1 for several traits, suggesting limited ability to capture additional environmental structure. In contrast, nonlinear kernels occasionally captured more complex patterns, leading to a slight reduction (e.g., DBH23 under soil) depending on the trait-covariate combination.

#### Reduction of residual variance

Overall, the inclusion of environmental covariates (W) and G×W interactions had limited effects on reducing residual variance (Figure 5 and Table 5). Models including W alone (M2) showed only minor and inconsistent changes relative to the baseline model, regardless of covariate type or kernel. Incorporating G×W interactions (M3) led to greater reductions in residual variance, with the largest improvements observed under the GK kernel (1.70% -2.78%), especially when soil covariates were fitted. However, reductions under the linear and AK kernels were generally less than 1.5%. Taken together, these results suggest that environmental covariates and reaction-norm terms explained only a small fraction of the residual variation not accounted for by the baseline model.

**Figure 5.**
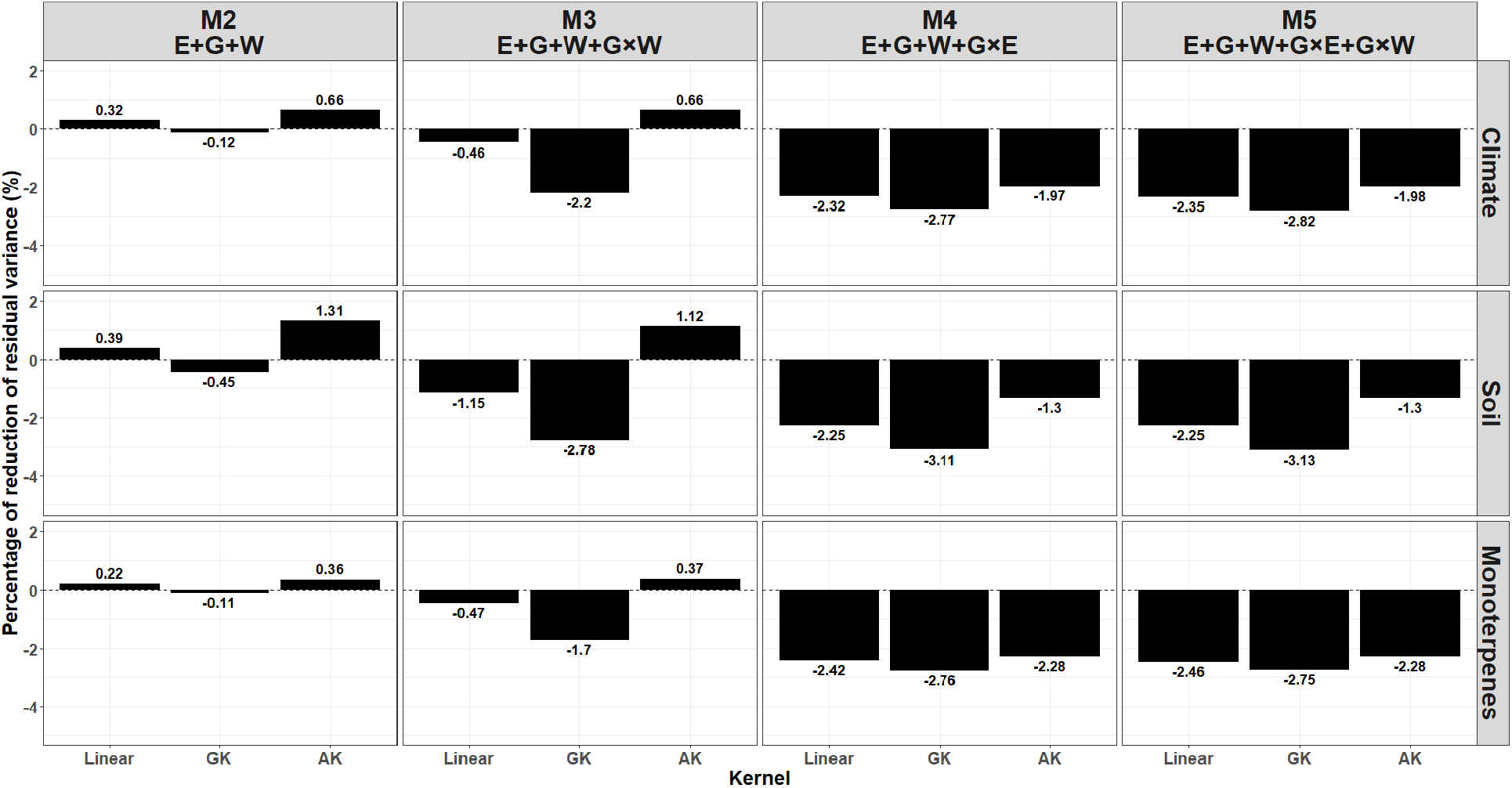
Averaged percentage reduction in residual (unexplained) variance relative to the baseline multi-environment ssGBLUP Model 1, which includes only environmental (site) and genetic effects (M1; E + G). Results are shown for Model 2 (M2; E + G + W), Model 3 (M3; E + G + W + G×W), Model 4 (M4; E + G + W + G×E), and Model 5 (M5; E + G + W + G×E + G×W), across traits, environmental covariates, and kernels approaches. Residual variances were obtained by averaging across traits. For nonlinear kernel approaches, results correspond to the Gaussian kernel (GK) with bandwidth *h* = 1 and the deep arc-cosine kernel (AK) with a single (1) layer.

**Table 5.** Average proportion of phenotypic variance explained by each variance component, and residual variance, across all studied traits for the fitted multi-environment ssGBLUP models. Models incorporated enviromic covariates (climatic and soil variables) and targeted metabolomic covariates (monoterpenes), modeled using a linear kernel (Linear) and nonlinear kernels, including the Gaussian kernel (GK; bandwidth parameter *h* = 1) and the deep arc-cosine kernel (AK; single hidden layer). Reported values represent averages across traits. See text for model abbreviations.

| **Model** | **Total explained phenotypic variance** |  |  | **Residual variance** |  |  |
|---|---|---|---|---|---|---|
|  | **Linear** | **GK** | **AK** | **Linear** | **GK** | **AK** |
| S+G | 13.92 |  |  | 86.08 |  |  |
| **Climate** |  |  |  |  |  |  |
| S+G+W | 13.64 | 14.02 | 13.35 | 86.36 | 85.98 | 86.65 |
| S+G+W+G×W | 14.32 | 15.82 | 13.35 | 85.68 | 84.18 | 86.65 |
| S+G+W+G×E | 15.92 | 16.31 | 15.62 | 84.08 | 83.69 | 84.38 |
| S+G+W+G×E+G×W | 15.95 | 16.34 | 15.62 | 84.05 | 83.66 | 84.38 |
| **Soil** |  |  |  |  |  |  |
| S+G+W | 13.59 | 14.31 | 12.80 | 86.41 | 85.69 | 87.20 |
| S+G+W+G×W | 14.92 | 16.31 | 12.96 | 85.08 | 83.69 | 87.04 |
| S+G+W+G×E | 15.86 | 16.60 | 15.04 | 84.14 | 83.40 | 84.96 |
| S+G+W+G×E+G×W | 15.86 | 16.61 | 15.04 | 84.14 | 83.39 | 84.96 |
| **Monoterpenes** |  |  |  |  |  |  |
| S+G+W | 13.73 | 14.01 | 13.62 | 86.27 | 85.99 | 86.38 |
| S+G+W+G×W | 14.33 | 15.39 | 13.60 | 85.67 | 84.61 | 86.40 |
| S+G+W+G×E | 16.00 | 16.29 | 15.89 | 84.00 | 83.71 | 84.11 |
| S+G+W+G×E+G×W | 16.04 | 16.29 | 15.89 | 83.96 | 83.71 | 84.11 |

When modeling the G×E interaction (M4), reductions in residual variance became more pronounced across all covariates. The linear kernels showed consistent moderate reduction (-2.25% to -2.42%). In contrast, nonlinear kernels provided slightly stronger reductions, particularly for the GK kernel (-2.76% to -3.11%), while the AK kernel showed similar but slightly smaller effects (-1.30% to -2.28%). These results indicate that a portion of the remaining unexplained variation is associated with G×E interactions not fully captured by W or G×W alone, although differences among kernels remain moderate.

When both interaction terms were included (M5: E + G + W + G×E + G×W), reductions in residual variance remained consistent across covariates and kernels but did not substantially exceed those observed for M4. The linear kernel showed stable reductions ranging from -2.25% to -2.46%, while the GK kernel again provided the largest effects (-2.75% to -3.13%), particularly for soil covariates. The AK kernel showed slightly smaller but comparable reductions (-1.30% to -2.28%). Across covariates, values were relatively homogeneous, ranging from -1.98% to -2.82% for climate, -1.30% to -3.13% for soil, and -2.28% to -2.75% for monoterpenes.

In summary, averaging across traits, differences among kernels were evident but depended on model complexity. Nonlinear approaches, particularly GK, generally outperformed the linear kernel, capturing a larger proportion of residual variance, whereas AK showed more variable performance and did not consistently improve over the linear approach in the most complex models. Moreover, the relative performance of covariates was stable, with soil variables generally providing the greatest reduction, followed by climate and monoterpenes.

#### Difference in explaining sources of variation

Consistent with this expectation, Figure 6 (and Table S3) shows that across the 12 traits, the additive genetic component (G) consistently explained the largest proportion of variance, averaging 8.14%, exceeding the combined environmental contribution of site and replication within site (4.10% and 0.58%, respectively). In the baseline model (M1; E + G), genetic variance accounted for 8.37% out of a total explained variance of 13.92%, indicating that the genomic component was the largest modeled source of variance. The inclusion of covariates in M2 resulted in modest changes in total explained variance (12.80-14.31%), with W explaining between 0.13% and 1.95% depending on the covariate type and kernel. Nonlinear kernels, particularly Gaussian (GK), consistently captured a greater proportion of variance. When genotype-by-covariate (G×W) interactions were included (M3), the G×W component explained an additional proportion of variance (mean = 0.60%, up to 2.94%), accompanied by a slight reduction in the additive genetic component (e.g., down to 7.15%). As previously mentioned, this suggests a redistribution of variance from G to interaction effects, indicating that part of the additive genetic signal is being captured by the G×W interaction component. Total explained variance increased up to 16.31%. The inclusion of G×E interaction (M4) resulted in an appreciable contribution of G×E (mean = 2.31%), increasing total explained variance to 15.04-16.60%. A slight reduction in G (8.02-8.13%) further indicates that part of the genetic variance is environment-dependent. Finally, the full model (M5), incorporating both G×E and G×W interactions, yielded the highest total explained variance (up to 16.61%). G×E remained the dominant interaction term, while G×W contributed more modestly (generally ≤0.50%). Nonlinear kernels, particularly GK, showed superior performance, explaining a slightly larger proportion of the variance (15.69%) than linear kernels (15.01%). Overall, total explained variance generally increased with model complexity from M1 to M5.

**Figure 6.**
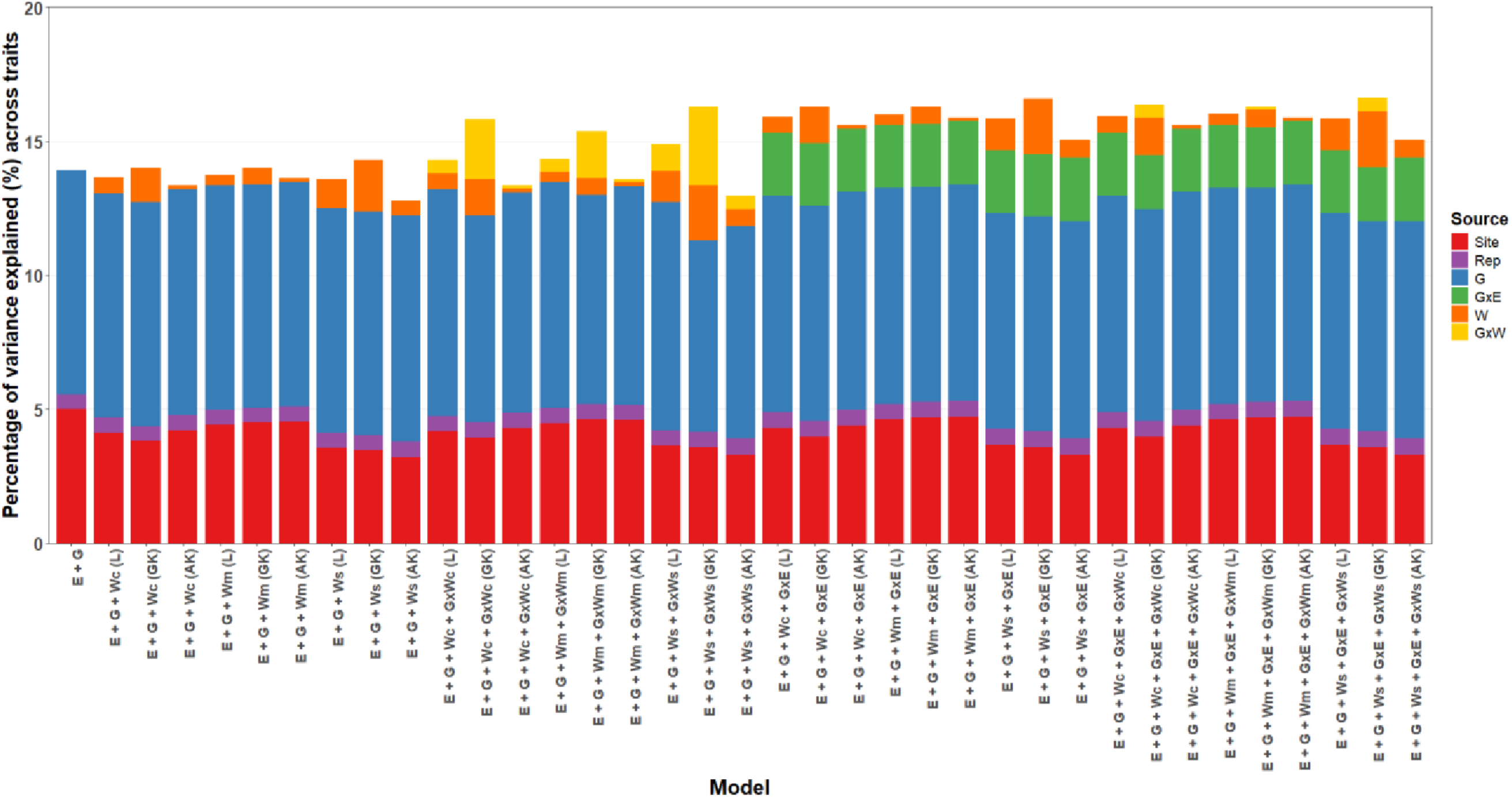
Variance component partitioning averaged across the twelve studied traits. Bars represent the percentage of variance explained by each source for each model averaged across trait including environment (Site), replication nested within site (Rep), additive genetic (G), genotype-by-environment interaction (G×E), environment covariates (W), and genotype-by-covariate interaction (G×W) components, as estimated using the multi-environment ssGBLUP mixed models. Results are shown for the five models studied. Models differ in the inclusion of climatic (c), soil (s), and targeted monoterpene (m) covariates, fitted using linear (L) and nonlinear kernels: Gaussian (GK; *h* = 1) and arc-cosine (AK; one layer). Model abbreviations are described in the text and Table 2.

This general increase in explained variance with model complexity was also evident at the trait level, although the relative importance of individual variance components varied among traits (Figure 7 and Table S2). Although G remained predominant, its importance differed substantially, ranging from high values in C13 (13.96-14.32%) and NSWGR (18.49-18.93%) to much lower contributions in Decline (mean = 3.01%) and early growth traits such as HT09 (2.44-4.99%). Height traits showed moderate G contributions in HT23 (5.79-7.25%) and HT30 (4.61-6.56%), increasing in HT35 (12.95-13.09%), while WD exhibited consistently high genetic contributions (9.05-11.77%). Environmental effects were relevant in early growth (HT09, up to 16.59%), remained moderate in HT23-HT35 (0.00-6.97%), and were negligible in NSWGR. The contribution of covariates (W) also varied widely: soil covariates explained substantial variance in HT09 (up to 20.60%) and DBH23 (up to 4.39%), while remaining negligible for NSWGR and HT30; climate covariates showed moderate effects in HT23 (up to 5.91%) and WD (up to 6.30%), while being minimal in HT30 and absent in NSWGR; monoterpenes showed generally limited and trait-dependent effects. Interaction components also showed strong heterogeneity. G×E was particularly relevant for Decline (up to 6.34%) and consistently important across height traits (HT09, HT23, HT30: 2.83-4.17%), as well as for WD (2.79-3.96%), but negligible in HT35 and NSWGR. Similarly, G×W interactions were trait-dependent, with notable contributions in Decline (up to 8.59%), WD (up to 6.02%), and HT09-HT30 (up to 4.21 and 4.95%, respectively), while remaining close to zero in HT35, NSWGR, and C13.

**Figure 7.**
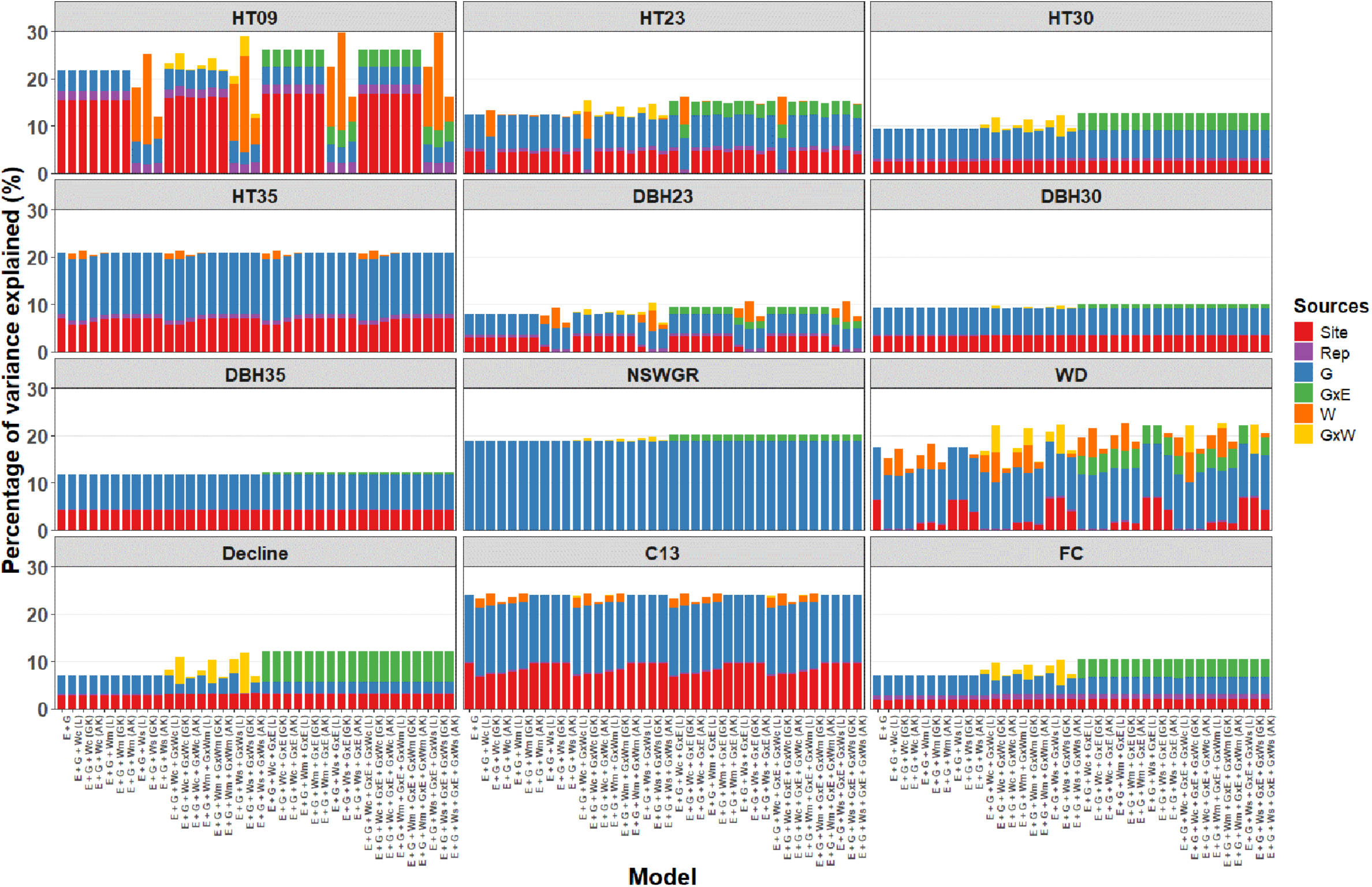
Variance component partitioning across twelve traits. Bars represent the percentage of variance explained by each source for each model, including environment (Site), replication nested within site (Rep), additive genetic (G), genotype-by-environment interaction (G×E), covariates (W), and genotype-by-covariate interaction (G×W) components, as estimated using the multi-environment ssGBLUP mixed models. Results are shown for the five models studied. Models differ in the inclusion of climatic (c), soil (s), and targeted monoterpene (m) covariates, fitted using linear (L) and nonlinear kernels: Gaussian (GK; *h* = 1) and arc-cosine (AK; one layer). Trait abbreviations are described in the text and Table S1.

### Predictive ability in untested environments

The Leave-One-Environment-Out cross-validation scheme revealed that the predictive performance of multi-environment ssGBLUP models in untested environments was generally high across traits and environments, with average predictive ability ranging from 0.688 to 0.701, depending on the model structure, covariate type, and kernel formulation (Figure 8 and Table S4). These results indicate that substantial predictive information could be transferred across environments, even under the baseline model (M1), suggesting moderate environmental connectivity among sites. Overall, predictive ability tended to increase with model complexity, although the magnitude of improvement varied considerably among environments, covariate types, and kernel formulations.

**Figure 8.**
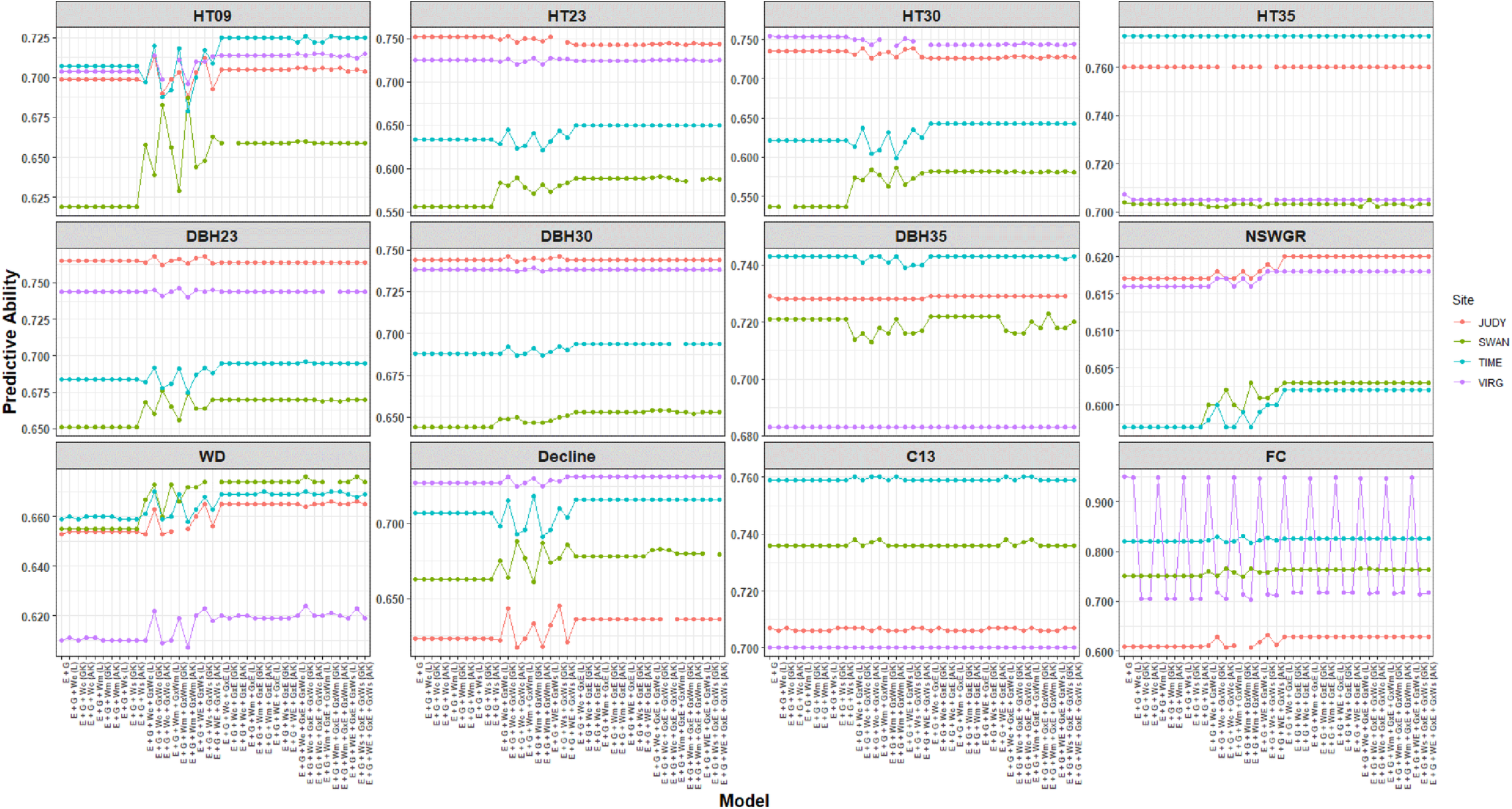
Predictive ability of multi-environment ssGBLUP reaction norm mixed models evaluated across twelve traits in each untested environment. Results are shown for the five models studied, covariate types (climate, soil and targeted monoterpenes), and kernels approaches (linear, Gaussian kernel and arc-cosine kernel). Trait and model abbreviations are described in the text and Table S1 and Table 2, respectively.

The incorporation of enviromic and targeted metabolomic covariates together with G×E and G×W interaction structures generally improved predictive ability relative to the baseline model, although the magnitude of improvement depended strongly on the environmental sensitivity of the trait and the target environment. Across all evaluated traits (Table 6 and Table S5), gains were moderate, typically ranging between 1 and 3% depending on the model and kernel. The largest improvements were consistently observed under models M4 and M5 (up to 3.65%), indicating that explicitly modeling genotype-by-environment heterogeneity improved the prediction of genotype performance in unobserved environments. Most of this gain appeared to be driven by the G×E component, whereas the additional contribution of G×W was comparatively small once G×E was included in the model. For example, under AK formulations with monoterpene covariates, predictive ability at SWAN increased from 0.653 under M1 to values close to 0.677 under M5. Overall, these results indicate that the benefits of increasing model complexity were generally positive but highly dependent on trait, environment, covariate type, and kernel formulation.

**Table 6.** Average Predictive ability across all studied traits for the fitted multi-environment ssGBLUP models. Models incorporated enviromic covariates (climatic and soil variables) and targeted metabolomic covariates (monoterpenes), modeled using a linear kernel (Linear) and nonlinear kernels, including the Gaussian kernel (GK; bandwidth parameter *h* = 1) and the deep arc-cosine kernel (AK; single hidden layer). Site and model abbreviations are described in the main text. See text for trait and enviroment abbrevations.

| **Model** | **Linear** |  |  |  | **GK** |  |  |  | **AK** |  |  |  |
|---|---|---|---|---|---|---|---|---|---|---|---|---|
|  | **JUDY** | **VIRG** | **SWAN** | **TIME** | **JUDY** | **VIRG** | **SWAN** | **TIME** | **JUDY** | **VIRG** | **SWAN** | **JUDY** |
| S+G | 0.700 | 0.721 | 0.653 | 0.699 |  |  |  |  |  |  |  |  |
| **Climate** |  |  |  |  |  |  |  |  |  |  |  |  |
| S+G+W | 0.699 | 0.721 | 0.653 | 0.700 | 0.700 | 0.701 | 0.663 | 0.699 | 0.699 | 0.701 | 0.653 | 0.700 |
| S+G+W+G×W | 0.699 | 0.722 | 0.666 | 0.697 | 0.706 | 0.704 | 0.662 | 0.706 | 0.691 | 0.699 | 0.671 | 0.694 |
| S+G+W+G×E | 0.702 | 0.722 | 0.669 | 0.708 | 0.702 | 0.703 | 0.670 | 0.708 | 0.702 | 0.703 | 0.669 | 0.708 |
| S+G+W+G×E+G×W | 0.703 | 0.723 | 0.670 | 0.708 | 0.703 | 0.703 | 0.670 | 0.708 | 0.709 | 0.703 | 0.669 | 0.708 |
| **Soil** |  |  |  |  |  |  |  |  |  |  |  |  |
| S+G+W | 0.699 | 0.721 | 0.653 | 0.699 | 0.700 | 0.701 | 0.653 | 0.699 | 0.700 | 0.701 | 0.653 | 0.699 |
| S+G+W+G×W | 0.702 | 0.723 | 0.663 | 0.698 | 0.696 | 0.703 | 0.665 | 0.705 | 0.698 | 0.699 | 0.669 | 0.701 |
| S+G+W+G×E | 0.702 | 0.722 | 0.669 | 0.708 | 0.702 | 0.703 | 0.669 | 0.708 | 0.702 | 0.703 | 0.669 | 0.708 |
| S+G+W+G×E+G×W | 0.702 | 0.722 | 0.669 | 0.708 | 0.703 | 0.703 | 0.668 | 0.708 | 0.700 | 0.703 | 0.669 | 0.708 |
| **Monoterpenes** |  |  |  |  |  |  |  |  |  |  |  |  |
| S+G+W | 0.699 | 0.721 | 0.653 | 0.700 | 0.699 | 0.701 | 0.653 | 0.700 | 0.699 | 0.701 | 0.653 | 0.700 |
| S+G+W+G×W | 0.699 | 0.722 | 0.666 | 0.696 | 0.714 | 0.699 | 0.658 | 0.705 | 0.697 | 0.698 | 0.672 | 0.692 |
| S+G+W+G×E | 0.702 | 0.722 | 0.669 | 0.708 | 0.702 | 0.703 | 0.669 | 0.708 | 0.702 | 0.703 | 0.669 | 0.708 |
| S+G+W+G×E+G×W | 0.703 | 0.723 | 0.669 | 0.709 | 0.702 | 0.699 | 0.668 | 0.708 | 0.703 | 0.703 | 0.677 | 0.708 |

Differences among covariate types were generally modest. Climatic and soil covariates produced relatively stable responses across kernels and environments, whereas targeted metabolomic covariates generated more variable responses, particularly under nonlinear kernels. Although linear kernels generally provided the highest and most stable average predictive performance across all traits and environments (Figure 9 and Table S6), nonlinear kernels occasionally produced larger gains in specific scenarios (Figure 8 and Table S4).

**Figure 9.**
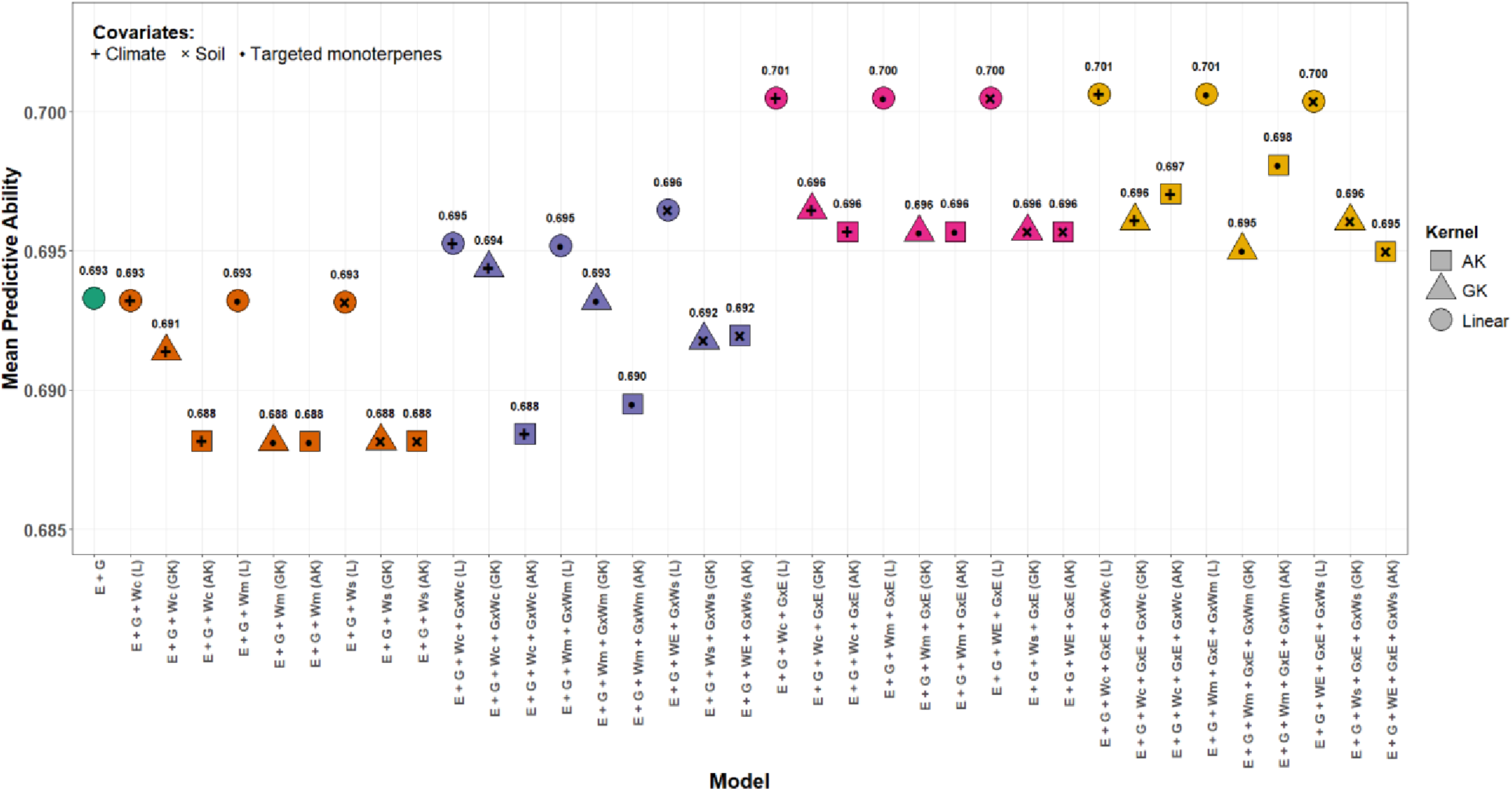
Predictive ability averaged across the twelve studied traits and four sites. Results are shown for the five models studied. Symbols indicate the type of covariates included: climatic (+), soil (×), and targeted monoterpenes (•), while shapes represent kernel approaches: linear, Gaussian kernel (GK; bandwidth *h* = 1), and arc-cosine kernel (AK; one layer). Numbers above symbols indicate the mean predictive ability for each model averaged across traits and sites.

Moreover, the advantages of incorporating covariates and interaction structures became substantially more evident for traits exhibiting stronger G×E patterns (average pedigree-based genetic correlations across site pairs < 0.65, Figure 2), namely HT09, HT23, HT30, and Decline (Table 7 and Table S7). For these traits, predictive ability improvements reached up to 5.81% under linear kernels, 5.94% under GK, and 7.85% under AK. For instance, under AK formulations with monoterpene covariates, predictive ability at SWAN increased from 0.594 under M1 to approximately 0.640 under M5. These gains were particularly evident in environmentally contrasting sites such as SWAN, where baseline predictive ability was comparatively low. In contrast, environments with already high baseline predictive performance showed comparatively smaller responses to model complexity.

**Table 7.**
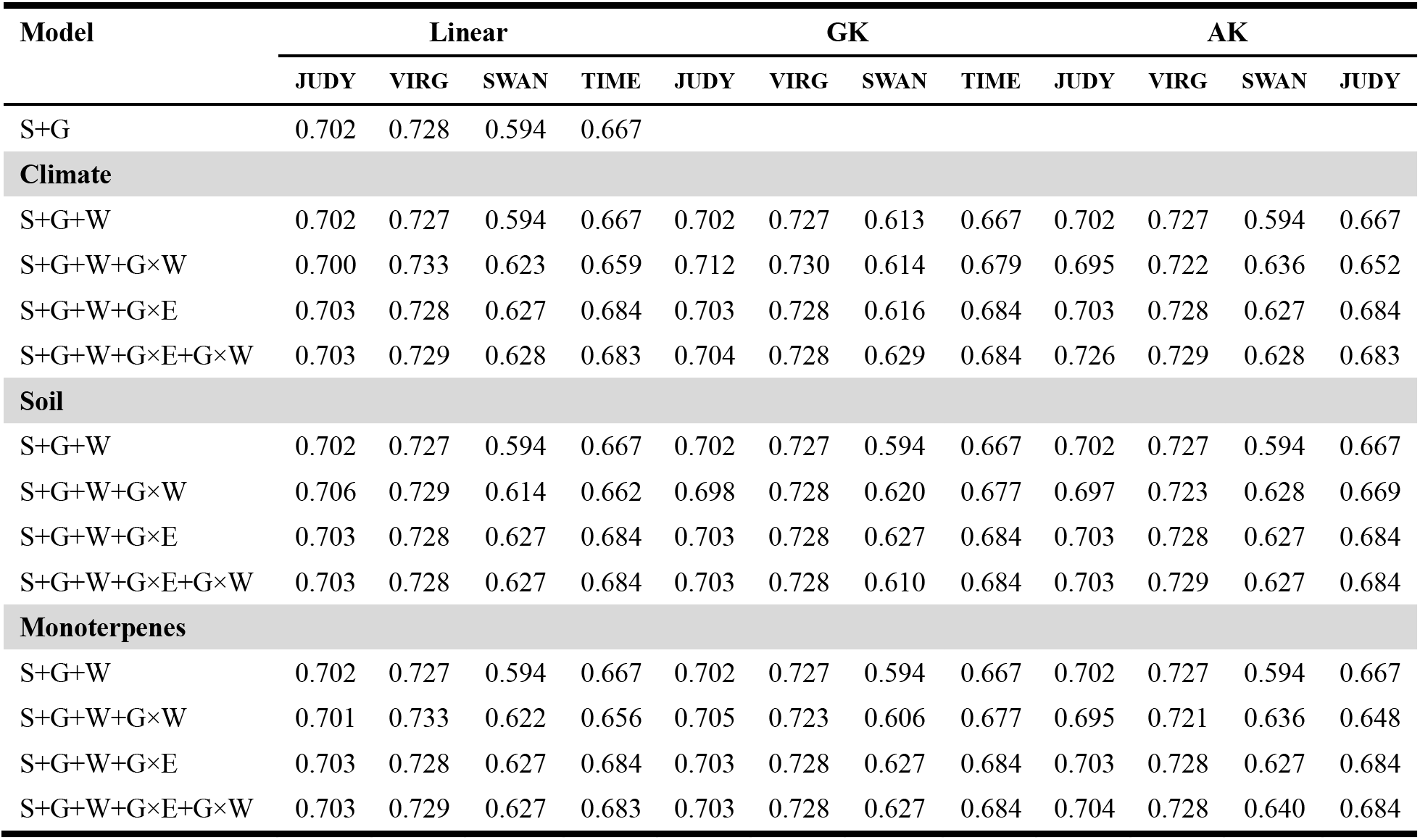
Average Predictive ability for the four traits showing the strongest G×E patterns (HT09, HT23, HT30, and Decline) (average pedigree-based genetic correlations across site pairs < 0.65, **Figure 2) across multi-environment ssGBLUP reaction-norm models.** Models incorporated enviromic covariates (climatic and soil variables) and targeted metabolomic covariates (monoterpenes), modeled using a linear kernel (Linear) and nonlinear kernels, including the Gaussian kernel (GK; bandwidth parameter *h* = 1) and the deep arc-cosine kernel (AK; single hidden layer). Site and model abbreviations are described in the main text. See text for trait and enviroment abbrevations.

| **Model** | **Linear** |  |  |  | **GK** |  |  |  | **AK** |  |  |  |
|---|---|---|---|---|---|---|---|---|---|---|---|---|
|  | **JUDY** | **VIRG** | **SWAN** | **TIME** | **JUDY** | **VIRG** | **SWAN** | **TIME** | **JUDY** | **VIRG** | **SWAN** | **JUDY** |
| S+G | 0.702 | 0.728 | 0.594 | 0.667 |  |  |  |  |  |  |  |  |
| **Climate** |  |  |  |  |  |  |  |  |  |  |  |  |
| S+G+W | 0.702 | 0.727 | 0.594 | 0.667 | 0.702 | 0.727 | 0.613 | 0.667 | 0.702 | 0.727 | 0.594 | 0.667 |
| S+G+W+G×W | 0.700 | 0.733 | 0.623 | 0.659 | 0.712 | 0.730 | 0.614 | 0.679 | 0.695 | 0.722 | 0.636 | 0.652 |
| S+G+W+G×E | 0.703 | 0.728 | 0.627 | 0.684 | 0.703 | 0.728 | 0.616 | 0.684 | 0.703 | 0.728 | 0.627 | 0.684 |
| S+G+W+G×E+G×W | 0.703 | 0.729 | 0.628 | 0.683 | 0.704 | 0.728 | 0.629 | 0.684 | 0.726 | 0.729 | 0.628 | 0.683 |
| **Soil** |  |  |  |  |  |  |  |  |  |  |  |  |
| S+G+W | 0.702 | 0.727 | 0.594 | 0.667 | 0.702 | 0.727 | 0.594 | 0.667 | 0.702 | 0.727 | 0.594 | 0.667 |
| S+G+W+G×W | 0.706 | 0.729 | 0.614 | 0.662 | 0.698 | 0.728 | 0.620 | 0.677 | 0.697 | 0.723 | 0.628 | 0.669 |
| S+G+W+G×E | 0.703 | 0.728 | 0.627 | 0.684 | 0.703 | 0.728 | 0.627 | 0.684 | 0.703 | 0.728 | 0.627 | 0.684 |
| S+G+W+G×E+G×W | 0.703 | 0.728 | 0.627 | 0.684 | 0.703 | 0.728 | 0.610 | 0.684 | 0.703 | 0.729 | 0.627 | 0.684 |
| **Monoterpenes** |  |  |  |  |  |  |  |  |  |  |  |  |
| S+G+W | 0.702 | 0.727 | 0.594 | 0.667 | 0.702 | 0.727 | 0.594 | 0.667 | 0.702 | 0.727 | 0.594 | 0.667 |
| S+G+W+G×W | 0.701 | 0.733 | 0.622 | 0.656 | 0.705 | 0.723 | 0.606 | 0.677 | 0.695 | 0.721 | 0.636 | 0.648 |
| S+G+W+G×E | 0.703 | 0.728 | 0.627 | 0.684 | 0.703 | 0.728 | 0.627 | 0.684 | 0.703 | 0.728 | 0.627 | 0.684 |
| S+G+W+G×E+G×W | 0.703 | 0.729 | 0.627 | 0.683 | 0.703 | 0.728 | 0.627 | 0.684 | 0.704 | 0.728 | 0.640 | 0.684 |

Overall, these results indicate that the incorporation of enviromic and targeted metabolomic covariates within multi-environment ssGBLUP models can improve predictive performance in untested environments, particularly for traits exhibiting stronger G×E interaction patterns. The linear kernel was the strongest general-purpose model, while GK and AK functioned as context-dependent alternatives, suggesting that their utility may depend on the degree of nonlinear genotype-by-environment relationships underlying the trait.

## Discussion

This empirical study evaluated, for the first time in a forest tree species, the integration of genomic, enviromic (climatic and soil), and metabolomic (targeted monoterpenes) information within multi-environment ssGBLUP reaction norm models using both linear and nonlinear kernels. Three major findings emerged. First, the benefits of incorporating environmental information depended strongly on the magnitude of G×E interaction, with the largest improvements observed for traits exhibiting lower genetic correlations among environments. Second, environmental covariates improved environmental characterization and facilitated prediction in untested environments, although most gains were attributed to the explicit modeling of G×E rather than to environmental covariates alone. Third, targeted monoterpene profiles achieved predictive performance comparable to that of climatic and soil covariates, suggesting that targeted metabolomic information can serve as a biologically meaningful descriptor of environmental similarity.

Previous studies evaluating genomic prediction (GP) across environments in forest trees have reported contrasting results. These results range from substantial declines in predictive ability when predictions are transferred to environmentally distinct sites (Resende *et al*., 2012b; Gamal El-Dien *et al*., 2015; Shalizi *et al*., 2022) to relatively stable performance when G×E interaction is limited (Beaulieu *et al*., 2014; Lenz *et al*., 2017), and even to improved prediction when G×E relationships are explicitly modeled through multi-environment kernels (Hu *et al*., 2023). This study demonstrates that the benefit of incorporating environmental information depends strongly on the magnitude of G×E interaction. Traits exhibiting lower genetic correlations among environments (<0.65; i.e., moderate G×E interaction; Figure S2), including HT09, HT23, HT30, and Decline, showed the largest improvements in model fit (Table 4), explained variance (Figure 7 and Table S2), and predictive ability (Figure 8 and Table S4) following the incorporation of G×E and environmental covariates. In contrast, traits characterized by high genetic correlations across sites, such as HT35 and DBH35, exhibited only marginal gains. These findings highlight that the value of reaction norm models increases as environmental sensitivity becomes more pronounced, providing greater insight into the environmental and biological factors underlying G×E interactions.

Historically, G×E in forest tree breeding has been modeled within multi-trait frameworks, in which environments are treated as discrete entities and genetic correlations are estimated between pairs of sites (Falconer and Mackay, 1996). While effective, this approach simplifies environmental variation into a limited set of site categories. From the reaction norm perspective, genotype performance is instead viewed as a continuous function across environmental gradients, with each environment representing a point along a continuum rather than a discrete category (Kirkpatrick and Heckman, 1989). By incorporating environmental covariates, reaction norm models characterize environments through continuous descriptors, allowing information to be shared across environmental gradients and facilitating prediction in environments that have not been directly observed (Resende *et al*., 2021). The relatively high predictive abilities obtained in the leave-one-environment-out validation suggest that substantial predictive performance could be transferred across sites, supporting this framework for prediction beyond directly observed trial environments. This capacity is particularly relevant as tree deployment environments may increasingly depart from the conditions represented in current trials.

The variance partitioning results further support this interpretation. Although the inclusion of environmental covariates reduced the proportion of variance attributed to the main site effect, indicating that climatic, soil, and targeted metabolomic descriptors captured part of the environmental heterogeneity among trials, the environmental similarity kernels generally exhibited broader distributions than the realized G×E correlation patterns observed among environments (Figure S1). This indicates that similarity in measured environmental conditions does not necessarily translate directly into similarity in genetic responses across sites. Nevertheless, despite providing an explicit characterization of site differences, G×W interactions accounted for less phenotypic variation than G×E interactions, suggesting that the measured covariates did not fully represent the environmental dimensions underlying genotype-specific responses among sites. This interpretation is consistent with the limitation noted by (Jarquín *et al*., 2014) and other authors (e.g., (Mumford *et al*., 2023)) for reaction norm models, namely that measured environmental covariates may not fully describe environmental differences among sites because some relevant environmental factors may not have been measured. Such a limitation could explain why G×E remained the dominant source of interaction variance in our analyses and produced the greatest reductions in residual variance (Figure 5 and Table 5). Moreover, the most complex models (M4 and M5) consistently achieved the highest levels of explained variance (Table 5). This shift in variance structure was most evident for environmentally sensitive traits (Figure 2; genetic correlation < 0.65), which showed a redistribution of explained variation away from additive genetic effects and toward G×E, G×W, site effects, and environmental covariates (Table S2). Despite the increased importance of environmental and interaction components for these traits, additive genetic effects remained the largest source of explained variation across most trait-model combinations (Figure 6 and Table S2).

Beyond these broad differences in variance partitioning, the specific environmental factors underlying site variation also differed among traits (Figure 7 and Table S2). For example, soil covariates almost completely absorbed the site effect for HT09 under both linear and nonlinear kernels, suggesting that differences in early growth among sites were predominantly driven by edaphic conditions rather than by macro-climatic variation. This pattern is consistent with the importance of soil properties and resource availability during the establishment phase, when juvenile trees are sensitive to local micro-environmental conditions (e.g., (Henneb *et al*., 2020)). Similar patterns have been reported in *Pinus radiata* D. Don (Li *et al*., 2015), where soil characteristics were associated with genetic and site-related variation at ages 1 and 6. Interestingly, no significant associations with these soil factors were detected at age 8, suggesting that the environmental drivers of genetic and site variation may change as trees develop. In contrast, climatic covariates completely absorbed the site effect for WD, suggesting that differences in wood density among sites were primarily driven by climatic conditions. The influence of climate on wood density has been extensively documented in forest tree species (Thomas *et al*., 2004; Nabais *et al*., 2018), with temperature and moisture conditions playing a key role in shaping wood formation and resulting wood properties. For instance, (Cabon *et al*., 2020) showed that temperature and water potential jointly regulate cambial activity and xylem development in conifers, thereby influencing wood properties across contrasting environments. Importantly, these differences in variance partitioning were also reflected in model predictive performance (Table 6 and Table 7). Across traits, particularly those exhibiting moderate to strong G×E interactions (Figure 2), the models accounting for a greater proportion of phenotypic variance generally achieved higher predictive ability (Figure S2), especially when additional variation was captured through G×E and G×W interaction terms. This finding suggests that improvements in predictive performance were associated with the representation of additional sources of genetic and environmental heterogeneity relevant to trait expression.

An interesting finding was that targeted monoterpene covariates achieved predictive performance comparable to climatic and soil descriptors, despite representing a fundamentally different source of information (Tables 6 and 7). Whereas climate and soil covariates characterize external environmental conditions, monoterpene profiles represent physiological and biochemical phenotypes that may integrate genetic effects with responses to the conditions experienced by individual trees. Their comparable predictive performance suggests that targeted monoterpene profiles contain information relevant to cross-environment prediction and may provide a biologically mediated complement to conventional environmental descriptors. Collectively, these findings suggest that targeted metabolomics can serve not only as an additional omics layer within genomic prediction frameworks, as demonstrated by (Guo *et al*., 2023), but also as a complementary representation of the physiological state associated with environmental and genetic differences among trees. These findings are consistent with recent studies highlighting the value of metabolomic information as an intermediate layer linking genotype and phenotype (Wang *et al*., 2023; Yoshioka *et al*., 2026). Because metabolite profiles integrate both genetic and environmental influences, they may capture biological variation that is not fully explained by additive genetic effects alone. In this context, the comparable predictive performance of monoterpenes and conventional environmental descriptors suggests that part of the environmental signal relevant for prediction may also be represented through the physiological responses of trees. Within reaction norm models, such information may help characterize genotype-specific responses along biologically relevant environmental gradients associated with G×E. However, the generally small contribution of G×W interactions, when G×E was also fitted, suggests that monoterpene profiles capture only a limited fraction of genotype-specific environmental variation beyond that already represented by the broader G×E term.

Although nonlinear kernels, particularly GK, generally improved model fit (Table S2), reduced residual variance, and explained a larger proportion of phenotypic variation (Figure 5 and Table 5), these advantages did not translate into higher predictive ability. Linear kernels instead generally produced the highest and most stable average predictive performance across traits and environments (Figure 9). This highlights an important distinction between explanatory and predictive performance. While nonlinear kernels may more effectively capture complex environmental and biological covariance structures (Montesinos-López *et al*., 2021), the additional complexity introduced by these models does not necessarily improve their ability to generalize to novel environments. Similar observations have been reported by (Lourenço *et al*., 2024), who showed that increasing model complexity does not automatically translate into superior prediction and that simpler linear approaches often remain competitive because of their robustness and reduced tuning requirements. More broadly, accumulating evidence suggests that the relative performance of machine-learning algorithms depends strongly on trait architecture and dataset characteristics, making it unlikely that any single method will consistently outperform all others across scenarios (Reinoso-Peláez *et al*., 2022; Lourenço *et al*., 2024).

The trait-dependent responses observed in our study are consistent with recent evidence suggesting that additional omics layers may provide complementary information to genomic data in some cases, although the magnitude of these benefits can vary substantially among traits (Montesinos-López *et al*., 2021). When genomic effects accounted for most of the explained variation, as observed for HT35 and C13 (Figure 7 and Table S2), the inclusion of more complex kernels and additional covariates yielded only marginal improvements. In contrast, environmentally sensitive traits characterized by stronger G×E interactions (Figure 2) benefited more from nonlinear kernels and metabolomic information (Table S5 and Table S7). Notably, the combination of targeted monoterpene covariates with the AK kernel produced the largest predictive gains, reaching up to 7.85% in environmentally contrasting sites such as SWAN. These findings suggest that the complementary information provided by metabolomic covariates may arise, at least in part, from their ability to capture environmental variation that is not fully represented by genomic data alone, rather than from providing substantial additional genetic information, as suggested by (Guo *et al*., 2025) in a metabolomic-genomic BLUP study. Such complementary environmental information may contribute to improved prediction across contrasting environments for environmentally sensitive traits.

An important aspect of this study is that prediction was evaluated under a leave-one-environment-out framework, which closely mimics the practical challenge of predicting performance of genetically connected material in environments not represented in the training population. Although future climatic conditions cannot be observed directly, the use of long-term historical weather records allowed environmental variability to be represented through continuous descriptors that capture both past environmental variation and cyclical climatic patterns, thereby introducing a temporally resolved dimension to GP (Howard and Lipka, 2025). Nevertheless, the relatively small number of environments evaluated in this study highlights the need for validation across broader environmental gradients before these approaches can be deployed for climate-adaptation strategies in forest tree breeding. Previous studies have reported larger predictive gains when reaction norm models are trained across a wider range of environments, suggesting that increasing both the number and diversity of test sites may further improve the characterization of environmental relatedness and genotype-specific responses (Resende *et al*., 2021).

Overall, our results demonstrate that the benefits of integrating environmental information and interaction terms into GP models depends strongly on the magnitude of the G×E interaction. Environmentally responsive traits benefited substantially from the incorporation of environmental descriptors and interaction terms, resulting in improved characterization of G×E patterns and enhanced prediction in untested environments. The comparable performance of metabolomic, climatic, and soil covariates further suggests that biologically informed omics layers can complement traditional enviromic approaches. Collectively, these findings demonstrate that integrating genomic, enviromic, and metabolomic information within multi-environment ssGBLUP reaction norm models provides a promising framework for improving prediction across heterogeneous and unobserved environments and informing climate-resilient breeding and deployment policies in forest tree species.

## Data availability statement

Genotyping-by-sequencing (GBS) raw reads used in this study have been deposited in NCBI SRA BioProject -PRJNA715165: https://www.ncbi.nlm.nih.gov/bioproject/715165. Information on the lodgepole pine trials, including pedigree and phenotypic data used and analyzed during the current study, will be made available in a GitHub repository upon acceptance of the manuscript.

## Acknowledgements

We would like to acknowledge the RES-FOR staff that collected and prepared the many lodgepole pine samples for this research: Laura Vehring, Pablo Chung, Jillian Dyck, Sarah Suzuk, Kristie Bui, Chris Arbter, Rob Johnstone, Jesse Shirton, Arial Eatherton, Calvin Jensen, and Michael Thomson. We also wish to thank all the technicians that work in the Thomaś, Erbilgiń and Mansfield labs. We also want to acknowledge the administrative coordination work of Stacy Bergheim. Finally, we thank Dr. Diego Jarquin for his insightful comments on the implementation and validation of reaction norm models.

## Author contributions

Eduardo P. Cappa: Conceptualization, Formal analysis, Investigation, Methodology, Software, Visualization, Writing -Original Draft, Writing -Review and Editing. Charles Chen: Data Curation, Resources, Writing -Review and Editing. Andy Benowicz: Data Curation, Resources, Funding acquisition, Writing -Review and Editing. Barb R. Thomas: Project Administration, Resources, Supervision, Validation, Writing -Review and Editing. Yousry A. El-Kassaby: Project Administration, Supervision, Validation, Writing -Review and Editing.

